# Stochastic character mapping of state-dependent diversification reveals the tempo of evolutionary decline in self-compatible Onagraceae lineages

**DOI:** 10.1101/210484

**Authors:** William A. Freyman, Sebastian Höhna

## Abstract

A major goal of evolutionary biology is to identify key evolutionary transitions that correspond with shifts in speciation and extinction rates. Stochastic character mapping has become the primary method used to infer the timing, nature, and number of character state transitions along the branches of a phylogeny. The method is widely employed for standard substitution models of character evolution. However, current approaches cannot be used for models that specifically test the association of character state transitions with shifts in diversification rates such as state-dependent speciation and extinction (SSE) models. Here we introduce a new stochastic character mapping algorithm that overcomes these limitations, and apply it to study mating system evolution over a time-calibrated phylogeny of the plant family Onagraceae. Utilizing a hidden state SSE model we tested the association of the loss of self-incompatibility with shifts in diversification rates. We found that self-compatible lineages have higher extinction rates and lower net-diversification rates compared to self-incompatible lineages. Furthermore, these results provide empirical evidence for the “senescing” diversification rates predicted in highly selfing lineages: our mapped character histories show that the loss of self-incompatibility is followed by a short-term spike in speciation rates, which declines after a time lag of several million years resulting in negative net-diversification. Lineages that have long been self-compatible, such as *Fuchsia* and *Clarkia*, are in a previously unrecognized and ongoing evolutionary decline. Our results demonstrate that stochastic character mapping of SSE models is a powerful tool for examining the timing and nature of both character state transitions and shifts in diversification rates over the phylogeny.

## 1 Introduction

Evolutionary biologists have long sought to identify key evolutionary transitions that drive the diversification of life (Szathmary and Smith 1995; Sanderson and Donoghue 1996). One method frequently used to test hypotheses about evolutionary transitions is stochastic character mapping on a phylogeny (Nielsen 2002; Huelsenbeck et al. 2003). While most ancestral state reconstruction methods estimate states only at the nodes of a phylogeny, stochastic character mapping explicitly infers the timing and nature of each evolutionary transition along the branches of a phylogeny. However, current approaches to stochastic character mapping have two major limitations: the commonly used rejection sampling approach proposed by Nielsen (2002) is inefficient for characters with large state spaces (Huelsenbeck et al. 2003; Hobolth and Stone 2009), and more importantly current methods only apply to models of character evolution that are finite state substitution processes. While the first limitation has been partially overcome through uniformization techniques (Rodrigue et al. 2008; Irvahn and Minin 2014; Landis et al. 2018), a novel approach is needed for models with infinite state spaces, such as models that specifically test the association of character state transitions with shifts in diversification rates. These models describe the joint evolution of both a character and the phylogeny itself, and define a class of widely used models called state-dependent speciation and extinction models (SSE models; Maddison et al. 2007; FitzJohn et al. 2009; FitzJohn 2010, 2012; Goldberg and Igić 2012; Magnuson-Ford and Otto 2012; Freyman and Höhna 2018).

In this work we introduce a method to sample character histories directly from their joint probability distribution, conditional on the observed tip data and the parameters of the model of character evolution. The method is applicable to standard finite state Markov processes of character evolution and also more complex models, such as SSE model, that are infinite state Markov processes. The method does not rely on rejection sampling and does not require complex data augmentation (Van Dyk and Meng 2001) schemes to handle unobserved speciation/extinction events. Our implementation directly simulates the number, type, and timing of diversification rate shifts and character state transitions on each branch of the phylogeny. Thus, when applying our method together with a Markov chain Monte Carlo (MCMC; Metropolis et al. 1953; Hastings 1970) algorithm we can sample efficiently from the posterior distribution of both character state transitions and shifts in diversification rates over the phylogeny.

To illustrate the usefulness of our method to sample stochastic character maps from SSE models, we applied the method to examine the association of diversification rate shifts with mating system transitions in the plant family Onagraceae. The majority of flowering plants are hermaphrodites, and the loss of self-incompatibility (SI), the genetic system that encourages outcrossing and prevents self-fertilization, is a common evolutionary transition (Stebbins 1974; Grant 1981; Barrett 2002). Independent transitions to self-compatibility (SC) have occurred repeatedly across the angiosperm phylogeny (Igic et al. 2008) and within Onagraceae (Raven 1979). Despite the repeated loss of SI, outcrossing is widespread and prevalent in plants, an observation that led Stebbins to hypothesize that selfing was an evolutionary dead-end (Stebbins 1957). Stebbins proposed that over evolutionary time selfing lineages will have higher extinction rates due to reduced genetic variation and an inability to adapt to changing conditions. However, Stebbins also speculated that selfing is maintained by providing a short-term advantage in the form of reproductive assurance. The ability of selfing lineages to self reproduce has long been understood to be potentially beneficial in droughts and other conditions where pollinators are rare (Darwin 1876), or after long distance dispersal when a single individual can establish a new population (Baker 1955).

Recent studies have reported higher net-diversification rates for SI lineages, supporting Stebbins’ deadend hypothesis (Ferrer and Good 2012; Goldberg et al. 2010; de Vos et al. 2014; Gamisch et al. 2015). Explicit phylogenetic tests for increased extinction rates in SC lineages have been performed in Solanceae (Goldberg et al. 2010), Primulaceae (de Vos et al. 2014), and Orchidaceae (Gamisch et al. 2015), and all of these studies reported lower overall rates of net-diversification in SC lineages compared to SI lineages. In these studies the association of mating system transitions with shifts in extinction and speciation rates was tested using the Binary State Speciation and Extinction model (BiSSE; Maddison et al. 2007). More recently, BiSSE has been shown to be prone to falsely identifying a positive association when diversification rate shifts are associated with another character *not* included in the model (Maddison and FitzJohn 2015; Rabosky and Goldberg 2015). One approach to reduce the possibility of falsely associating a character with diversification rate heterogeneity is to incorporate a second, unobserved character into the model (i.e., a Hidden State Speciation and Extinction (HiSSE) model; Beaulieu and OMeara 2016; Caetano et al. 2018). The changes in the unobserved character’s state represent background diversification rate changes that are not correlated with the observed character. Our work here is the first to apply a HiSSE-type model to test Stebbins’ dead-end hypothesis. We additionally use simulations and Bayes factors (Kass and Raftery 1995) to evaluate the false positive error rate of our model. Most notably, we employ our novel stochastic character mapping method to reconstruct the timing of both diversification rate shifts and transitions in mating system over a fossil-calibrated phylogeny of Onagraceae. We test the hypothesis that SC lineages have higher extinction and speciation rates yet lower net-diversification rates compared to SI lineages, and investigate the short-term versus long-term macroevolutionary consequences of the loss of SI.

## 2 Methods

### 2.1 Stochastic Character Mapping Method

Figure 1 gives a side by side comparison of the standard stochastic character mapping algorithm as originally described by Nielsen (2002) and the approach introduced in this work. In standard stochastic character mapping the first step is to traverse the tree post-order (tip to root) calculating the conditional likelihood of the character being in each state at each node using Felsenstein’s pruning algorithm (Figure 1a; Felsenstein 1981). Transition probabilities are computed along each branch using matrix exponentiation. Ancestral states are then sampled at each node during a pre-order (root to tip) traversal (Figure 1b). Finally, character histories are repeatedly simulated using rejection sampling for each branch of the tree (Figure 1c).

A detailed pseudocode formulation of our new stochastic character mapping algorithm is provided in Algorithm 1. In this algorithm we begin similarly by traversing the tree post-order and calculating conditional likelihoods. However, instead of using matrix exponentiation we calculate the likelihood using a set of differential equations similar to Maddison et al. (2007). We numerically integrate these equations for every arbitrarily small time interval along each branch, however, unlike Maddison et al. (2007), we store a vector of conditional likelihoods for the character being in each state for every small time interval (Figure 1e). Letting 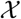 represent the observed tip data, Ψ an observed phylogeny, and *θ_q_* a particular set of character evolution model parameters, the likelihood at the root of the tree is then given by:

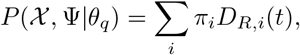

where *π_i_* is the root frequency of state *i* and *D_R,i_*(*t*) is the likelihood of the root node being in state *i* conditional on having given rise to the observed tree Ψ and the observed tip data 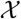 (Freyman and Höhna 2018).

We then sample a complete character history during a pre-order tree traversal. First, the root state is drawn from probabilities proportional to the marginal likelihood of each state at the root 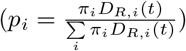. Then, states are drawn for each small time interval moving towards the tip of the tree conditioned on the state of the previous small time interval (Figure 1f). To compute the probability of a state at the end of each small time interval, we integrate numerically over a set of differential equations during this root-to-tip tree traversal, see Figure 1f. This integration, however, is performed in forward-time, thus a different and new set of differential equations must be used (defined below). With this approach we can directly sample character histories from a SSE process in forward-time, resulting in a complete stochastic character map sample without the need for rejection sampling or uniformization, see Figure 1.

### 2.2 Derivation of our differential equations

The two functions we integrate numerically are *D_N,i_*(*t*), which is defined as the probability that a lineage in state *i* at time *t* evolves into the observed clade *N*, and *E_i_*(*t*) which is the probability that a lineage in state *i* at time *t* goes extinct before the present, or is not sampled at the present.

In the following section we will derive the differential equations for our algorithm to compute the probability of the observed lineages and the extinction probabilities both backwards and forwards in time. We additionally show how the forward-time equations must be modified to handle non-reversible models of character evolution when sampling ancestral states or stochastic character maps.

**Figure 1:**
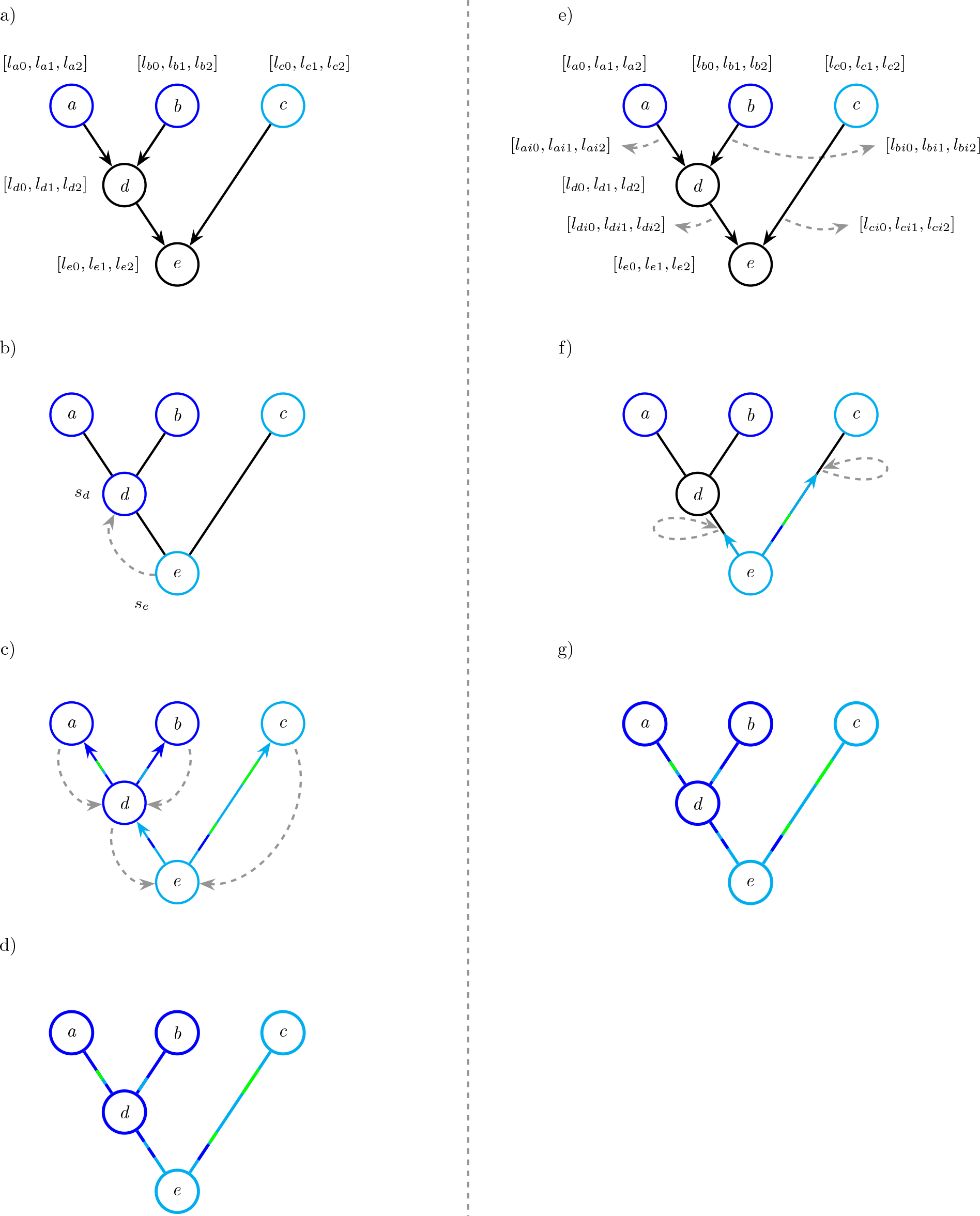
Schematic and comparison of stochastic character mapping methods. On the left *(a, b, c, d)* is an illustration of the standard stochastic character mapping algorithm as originally described by Nielsen (2002). On the right *(e, f, g)* is the approach introduced in this work. The first step in standard stochastic character mapping is *(a)* traversing the tree post-order (tip to root) calculating conditional likelihoods for each node. Next, ancestral states are sampled at each node during a pre-order (root to tip) traversal *(b)*. Branch by branch, character histories are then repeatedly simulated using rejection sampling *(c)*, resulting in a full character history *(d)*. The first step in the stochastic character mapping method introduced in this work is *(e)* traversing the tree post-order calculating conditional likelihoods for every arbitrarily small time interval along each branch and at nodes. Here the vector [*l*_*n*0_,*l*_*n*1_,*l*_*n*2_] represents the conditional likelihoods of the process at node n in states 0, 1, and 2, and the vector [*l*_*ni*0_,*l*_*ni*1_,*l*_*ni*2_] represents the conditional likelihoods of the process in the small time interval *i* along the branch leading to node n. Next, during a pre-order traversal ancestral states are sampled for each time interval *(f)*. The grey dashed loop represents the forward-time equations (Equations 4 and 6) conditioning on a state sampled during each small time interval. The result is a full character history *(g)* without the need for a rejection sampling step. See the main text for more details.

#### Algorithm 1

Stochastic character mapping algorithm. *D_Ni_*(*t*) is the probability that a lineage in state *i* at time *t* evolves into the observed clade *N*. *E_i_*(*t*) is the probability that a lineage in state *i* at time *t* goes extinct or is not sampled before the present.

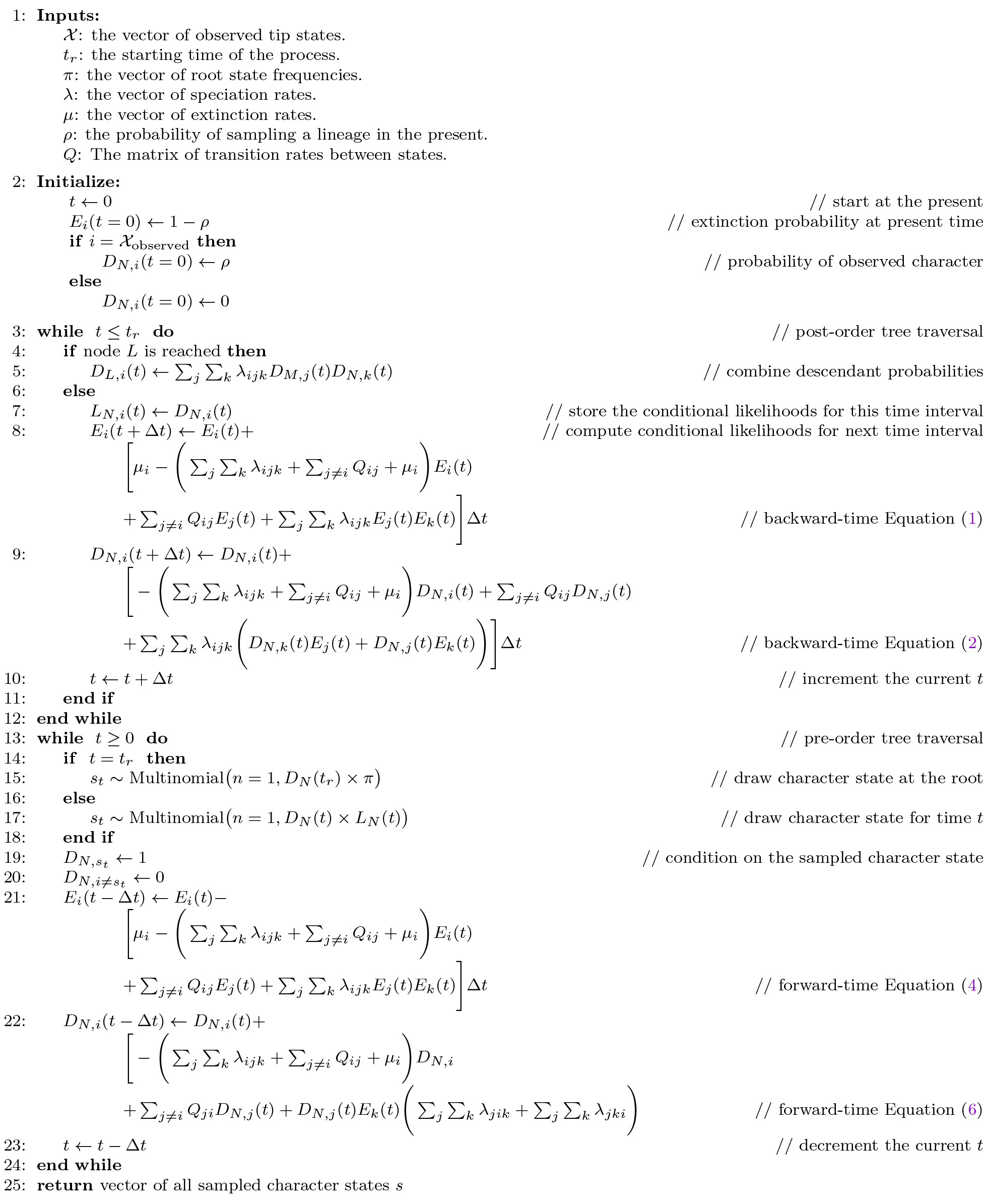

**Figure 2:**
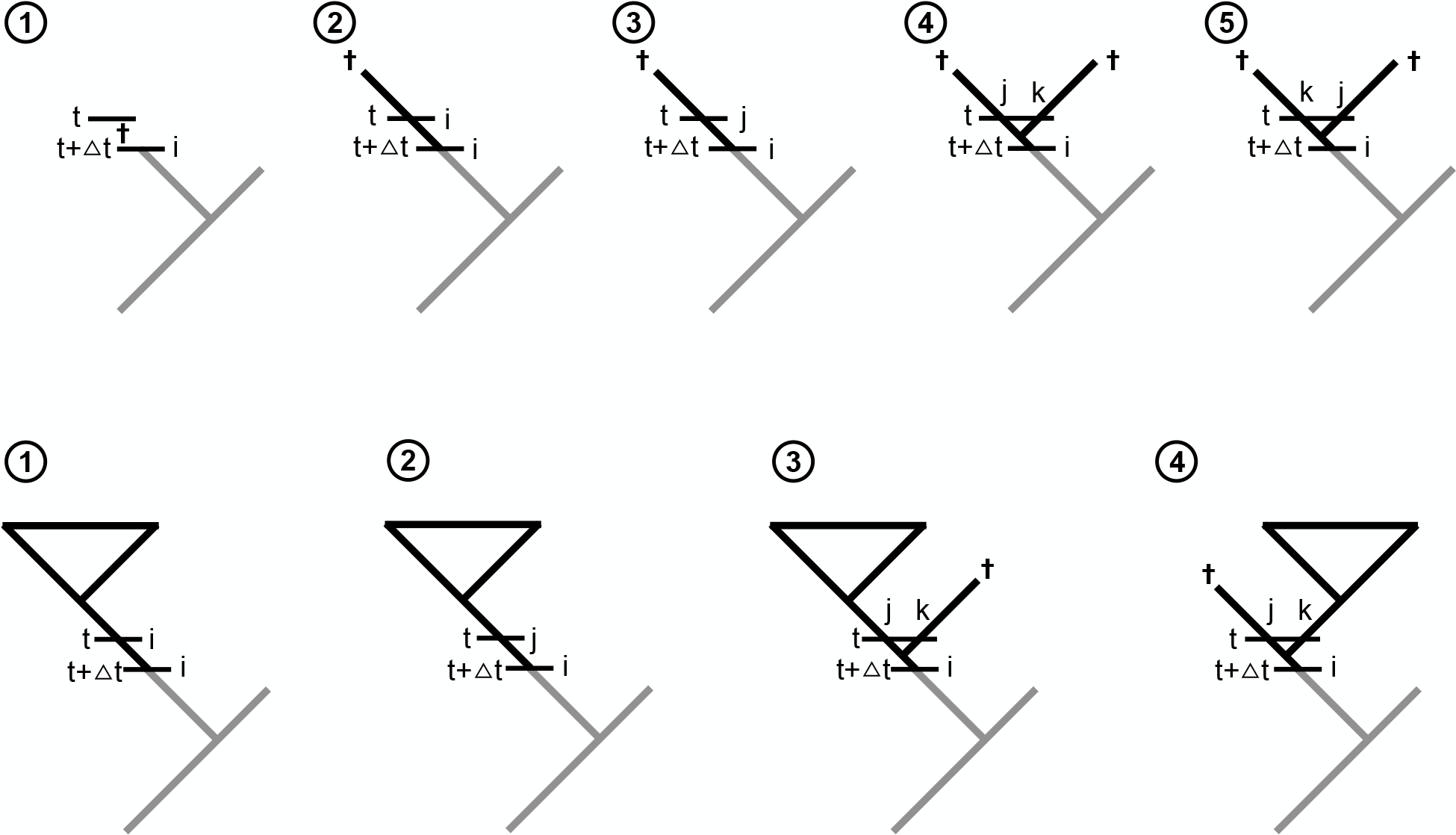
Alternative scenarios of events in a small time interval Δ*t* looking backwards in time. The top row shows the different scenarios for a lineage that goes extinct before the present. Case 1: The lineage goes extinct in the time interval Δ*t*. Case 2: There is no event in the time interval Δ*t* and the lineage goes extinct before the present. Case 3: The lineage undergoes a state-shift event to state *j* in the time interval Δ*t* and the lineage goes extinct before the present. Case 4: The lineage speciates and leaves a left daughter lineage in state *j* and a right daughter lineage in state *k* and both daughter lineages go extinct before the present. Case 5: The lineage speciates and leaves a left daughter lineage in state *k* and a right daughter lineage in state *j* and both daughter lineages go extinct before the present. The bottom row shows the different scenarios for an observed lineage. Case 1: There is no event in the time interval Δ*t*. Case 2: The lineage undergoes a state-shift event to state *j* in the time interval Δ*t*. Case 3: The lineage speciates and leaves a left daughter lineage in state *j* and a right daughter lineage in state *k* and only the left daughter lineage survives. Case 4: The lineage speciates and leaves a left daughter lineage in state *j* and a right daughter lineage in state *k* and only the right daughter lineage survives.

#### 2.2.1 Differential equations backwards in time

The original derivation of the differential equations for the state-dependent speciation and extinction (SSE) process are defined backward in time (Maddison et al. 2007). Here we use a generalization of the SSE process to allow for cladogenetic events where daughter lineages may inherit different states, as derived by Goldberg and Igić (2012), see also Magnuson-Ford and Otto (2012) and Ng and Smith (2014). We repeat this known derivation of the backwards process to show the similarities to our forward in time derivation. We present an overview of the possible scenarios of what can happen in a small time interval δ*t* in Figure 2. We need to consider all these scenarios in our differential equations.

First, let us start with the computation of the extinction probability. That is, we want to compute the probability of a lineage going extinct at time *t* + Δ*t*, denoted by *E*(*t* + Δ*t*), before the present time *t* = 0. We assume that we know the extinction probability of a lineage at time *t*, denoted by *E*(*t*), which is provided by our initial condition that *E*(*t* = 0) =0 because the probability of a lineage alive at the present cannot go extinct before the present, or *E*(*t* = 0) = 1 − *ρ* in the case of incomplete taxon sampling. We have five different cases (top row in Figure 2): (1) the lineage goes extinct within the interval Δ*t*; (2) nothing happens in the interval Δ*t* but the lineage eventually goes extinct before the present; (3) a state-change to state *j* occurs and the lineage now in state *j* goes extinct before the present; (4) the lineage speciates, giving birth to a left daughter lineage in state *j* and a right daughter lineage in state *k* and both lineages eventually go extinct before the present, or; (5) the lineage speciates, giving birth to a left daughter lineage in state *k* and a right daughter lineage in state *j* and both lineages eventually go extinct before the present. With this description of all possible scenarios we can derive the differential equation.

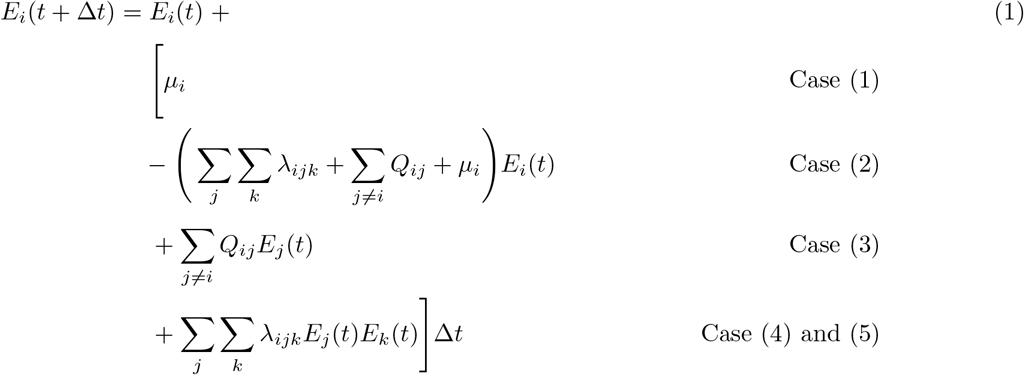

Similarly, we can consider all possible scenarios for an observed lineage. We have four different cases (bottom row in Figure 2): (1) nothing happens in the interval Δ*t*; (2) a state-change to state *j* occurs; (3) the lineage speciates, giving birth to a left daughter lineage in state *j* and a right daughter lineage in state *k* and only the left daughter lineage survives until the present, or; (4) the lineage speciates, giving birth to a left daughter lineage in state *j* and a right daughter lineage in state *k* and only the right daughter lineage survives until the present. Again, these scenarios are sufficient to derive the differential equation for the probability of an observed lineage, denoted *D*(*t*).

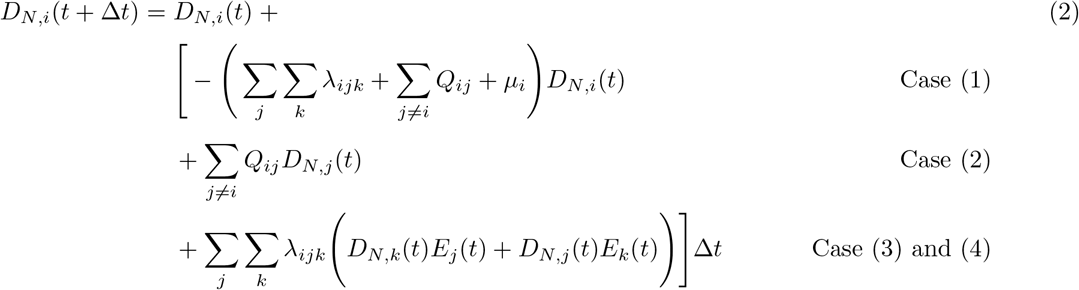

#### 2.2.2 Differential equations forward in time

Next, we want to compute the probability of extinction and the probability of an observed lineage forward in time. For the probability of extinction this is, in principle, almost identical to the backward in time equations. However, now we assume that we know *E*(*t*) and want to compute *E*(*t* − Δ*t*). We already computed *E*(*t*_*root*_) and *D*(*t*_*root*_) in our post-order tree traversal (from the tips to root). We use *E*(*t*_*root*_) as the initial conditions to approximate *E*(*t* − Δ*t*). Again, we have the same five different cases (top row in Figure 2): (1) the lineage goes extinct within the interval Δ*t*; (2) nothing happens in the interval Δ*t* but the lineage eventually goes extinct before the present; (3) a state-change to state *j* occurs and the lineage now in state *j* goes extinct before the present; (4) the lineage speciates, giving birth to a left daughter lineage in state *j* and a right daughter lineage in state *k* and both lineages eventually go extinct before the present, or; (5) the lineage speciates, giving birth to a left daughter lineage in state *k* and a right daughter lineage in state *j* and both lineages eventually go extinct before the present. However, these are the events that can happen in the future and we included the probabilities of these events already in *E*(*t*). Thus, we need to subtract instead of adding all possible scenarios that lead to the extinction of the lineage in the time interval Δ*t* from *E*(*t*) to obtain *E*(*t* − Δ*t*). This gives us the differential equation for the extinction probability as

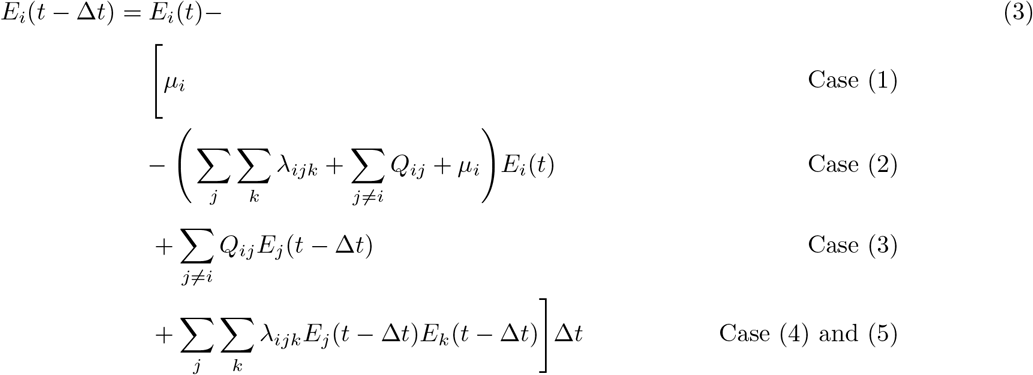

Unfortunately, we cannot solve Equation (3) directly because we do not know *E_j_*(*t* − Δ*t*) and *E_k_*(*t* − Δ*t*). Instead, we will approximate Equation (3) by using *E_j_*(*t*) instead of *E_j_*(*t* − Δ*t*), and *E_k_*(*t*) instead of *E_k_*(*t* − Δ*t*), respectively. Our approximation yields the new differential equation of the extinction probability by

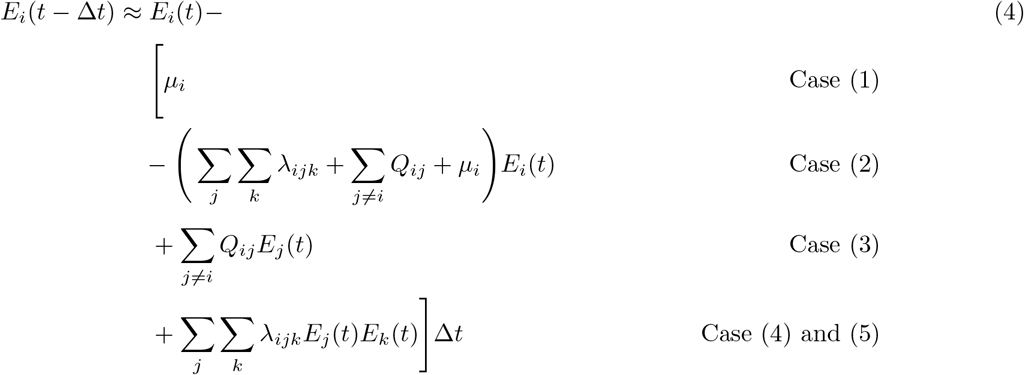

The derivation of the probability of an observed lineage in forward time is slightly different. When sampling a character history from the process we must compute *D*(*t* − Δ*t*) conditioned upon the character state sampled at time *t*. This does not effect the probability of a lineage going extinct before the present, so we can use *E*(*t*_*root*_) as the initial conditions to approximate *E*(*t* − Δ*t*). The initial conditions for the probability of an observed lineage, on the other hand, must account for the sampled character state. For example, if we sample the state *a* at time *t* our initial conditions to compute *D*(*t* − Δ*t*) must be *D_a_*(*t*) = 1.0 and *D_b_*(*t*) = 0.0 for all other character states *b*. Additionally, we must consider the process in forward time with all possible scenarios instead of backwards in time and subtracting the possible scenarios. We have four different cases that are similar to the cases for the backward in time computation (bottom row in Figure 2), however here the character state transitions are reversed since we are looking forward in time: (1) nothing happens in the interval Δ*t*; (2) with probability *D_N,j_*(*t*) the lineage was in state *j* and then a state-change to state *i* occurs; (3) with probability *D_N,j_*(*t*) the lineage was in state *j* and then speciates, giving birth to a left daughter lineage in state *i* and a right daughter lineage in state *k* and only the left daughter lineage survives until the present (the probability of extinction of the right daughter lineage is given by *E*_*k*_ (*t* − Δ*t*)), or; (4) with probability *D_N,j_*(*t*) the lineage was in state *j* and then speciates, giving birth to a left daughter lineage in state *k* and a right daughter lineage in state *i* and only the right daughter lineage survives until the present (the probability of extinction of the left daughter lineage is given by *E_k_*(*t* − Δ*t*)). From these four scenarios we derive the differential equation.

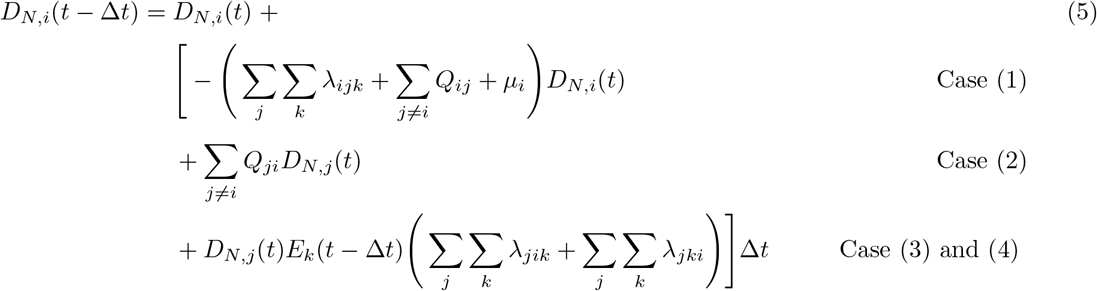

As before, we cannot solve Equation (5) directly because we do not know *E_k_*(*t* − Δ*t*). Thus, we use the same approximation as before and substitute *E_k_*(*t*) for *E_k_*(*t* − Δ*t*). This substitution gives our approximated differential equation.

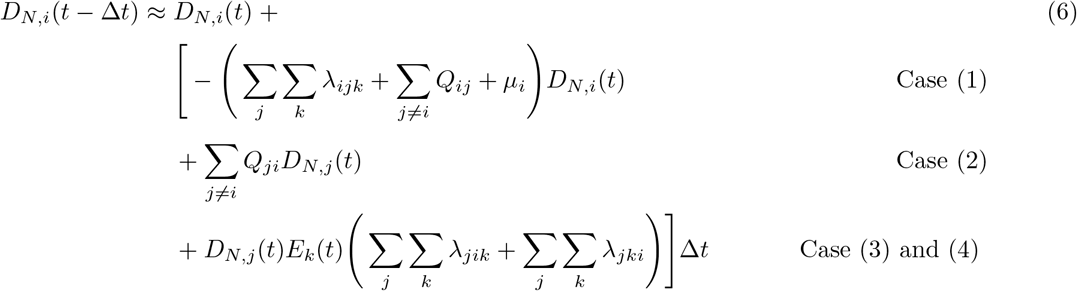

To sample character histories from an SSE process in forward-time during Algorithm (1) we calculate *E*(*t* − Δ*t*) using the approximation given by Equation (4) and *D*(*t* − Δ*t*) using Equation (6).

### 2.3 Correctness of the forward time equations

#### 2.3.1 Validation of the forward time extinction probabilities

For the purpose of demonstrating our forward time equations, we will use a non-symmetrical BiSSE model with states 0 and 1 which have the speciation rates λ_0_ = 1 and λ_1_ = 2, the extinction rates *μ*_0_ = 0.5 and *μ*_1_ = 1.5, and the transition rates *Q*_01_ = 0.2 and *Q*_10_ = 2.0. For simplicity we assume that there are no state changes at speciation events. We will first show that the approximations given by Equation (4) actually converge to the true probability of extinction if the time interval Δ*t* is very small (goes to zero). Note that we cannot show the same behavior for the forward in time probabilities of the observed lineage, *D*(*t*), because when conditioning on a sampled character state the forward in time probabilities will be different than the backward in time probabilities. For these probabilities we provide a different type of validation in Section 2.3.2.

**Figure 3:**
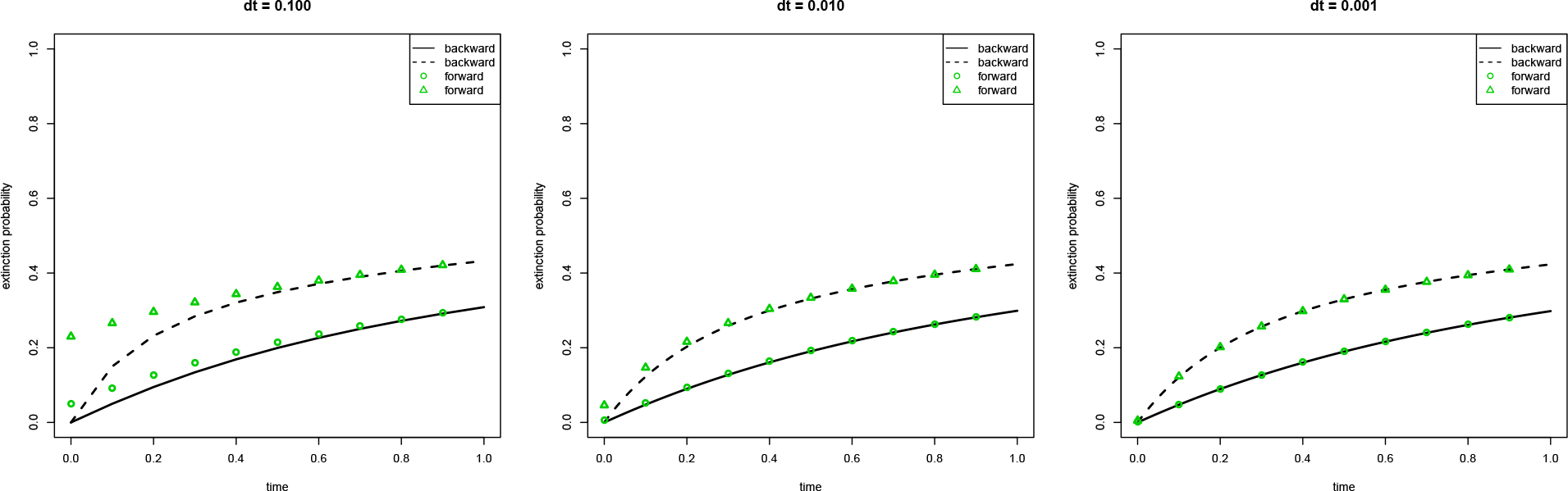
The probability of extinction computed backward and forward in time. Here we compute the extinction probabilities *E*_0_(*t*) and *E*_1_(*t*) for a BiSSE model backward and forward in time. Details about the parameters of the BiSSE model are given in the text. We varied the step-size Δ*t* for the numerical integration between 0.1, 0.01, and 0.001 to show that both computations give the same probabilities once Δ*t* is small enough.

We start by computing the probability of extinction and the probability of an observed lineage backward in time for a total time interval of 1.0. We initialize the computation with *E_i_*(*t* = 0) =0 and then compute *E*_0_(*t*) and *E*_1_(*t*) backward in time. Then, we use the computed values of *E_i_*(*t* = 1) as the initial values for our forward in time computation. If our approximation is correct, then we should get identical values for the extinction probabilities *E_i_*(*t*) for any value of *t*.

Figure 3 shows our computation using three different values for Δ*t*: 0.1, 0.01 and 0.001. We observe that our approximation of the forward in time computation of the probabilities converges to the backward in time computation when Δ*t* ≤ 0.001, which confirms our expectation. An explanation for the convergence is that *E*_0_(*t*) will be approximately equal to *E*_0_(*t* − Δ*t*), (and *E*_1_(*t*) to *E*_1_(*t* − Δ*t*)) the smaller Δ*t* becomes. In our actual implementation in `RevBayes` we use an initial step-size of Δ*t* = 10^−7^ but apply an adaptive numerical integration routine to minimize the error in the integrated function.

#### 2.3.2 Validation of the forward time equations against diversitree

**Figure 4:**
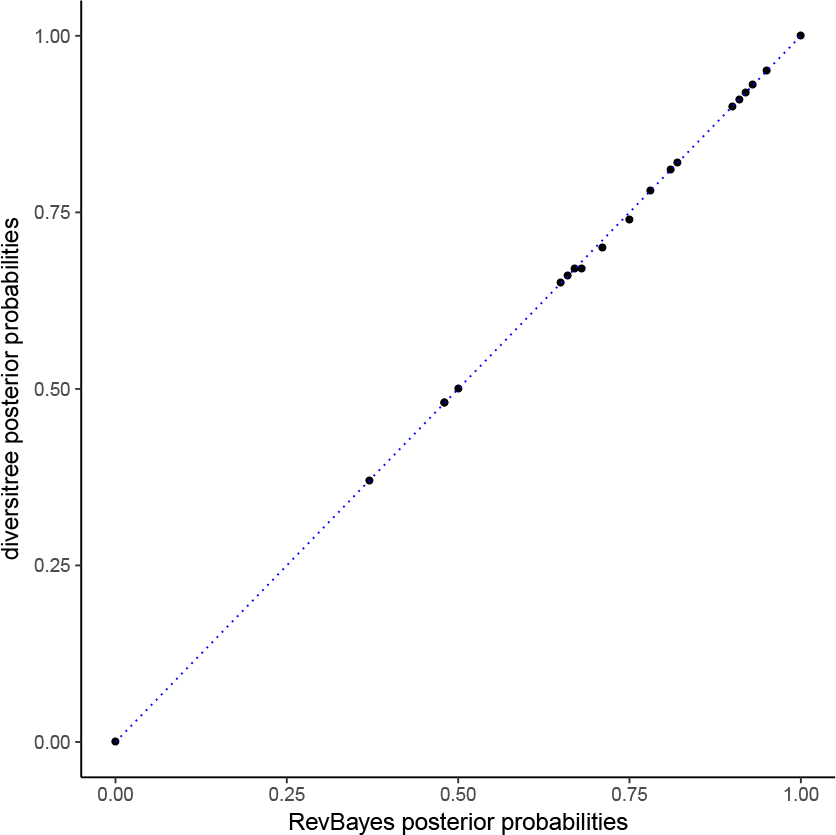
Comparing marginal posterior ancestral state estimates from diversitree to those calculated in RevBayes. Each point represents the posterior probability of a given node having the ancestral state 0. On the y-axis are the posterior probabilities as analytically calculated by `diversitree`. On the x-axis are the posterior probabilities as calculated by RevBayes using Algorithm (1). Our approximation given in Equation (6) yields highly similar posterior probabilities of the ancestral states as `diversitree`. Scripts to repeat this test with various parameter settings are provided in https://github.com/wf8/anc_state_validation.

Second, we validate our method of sampling character histories from an SSE process in forward-time by testing it against the analytical marginal ancestral state estimation implemented in the `R` package `diversitree` (FitzJohn 2012). Our method as implemented in `RevBayes` works for sampling both ancestral states and stochastic character maps, however `diversitree` can not sample stochastic character maps. Thus we limit our comparison to ancestral states estimated at the nodes of a phylogeny. Though our method works for all SSE models nested within ClaSSE, ancestral state estimation for ClaSSE is not implemented in `diversitree`, so we further limit our comparison to ancestral state estimates for a BiSSE model. Note that as implemented in `RevBayes` the BiSSE, ClaSSE, MuSSE (FitzJohn 2012), HiSSE (Beaulieu and OMeara 2016), ChromoSSE (Freyman and Höhna 2018), and GeoSSE (Goldberg et al. 2011) models use the same C++ classes and algorithms for parameter and ancestral state estimation, so validating under BiSSE should provide confidence in estimates made by `RevBayes` for all these SSE models.

Our method samples character histories from SSE models from their joint distribution conditioned on the tip states and the model parameters during MCMC. In contrast, `diversitree` computes marginal ancestral states analytically. Thus to directly compare results from these two approaches we calculated the marginal posterior probability of each node being in each state from a set of 10000 samples drawn by our Monte Carlo method. Figure 4 compares these estimates under a non-reversible BiSSE model where the tree and tip data were simulated in `diversitree` with the following parameters: λ_0_ = 0.2, λ_1_ = 0.4, *μ*_0_ = 0.01, *μ*_1_ = 0.1, and *q*_01_ = 0.1, *q*_10_ = 0.4. Figure 4 shows that using the approximation of *E*(*t* − Δ*t*) given by Equation (4) and the approximation to compute *D*(*t* − Δ*t*) in Equation (6) during Algorithm (1) results in marginal posterior estimates for the ancestral states that are nearly identical (up to some expected numerical and sampling errors) to those calculated analytically by `diversitree`. Scripts to perform this test with various parameter settings are provided in https://github.com/wf8/anc_state_validation.

### 2.4 Implementation, MCMC Sampling and Computation Efficiency

The stochastic character mapping method described here is implemented in C++ in the software `RevBayes` (Höhna et al. 2014, 2016). The RevGadgets R package (available at https://github.com/revbayes/RevGadgets) can be used to generate plots from `RevBayes` output. Scripts to run all `RevBayes` analyses presented here can be found in the repository at https://github.com/wf8/onagraceae.

Our method approximates the posterior distribution of the timing and nature of all character transitions and diversification rate shifts by sampling a large number of stochastically mapped character histories using MCMC. Uncertainty in the phylogeny and other parameters is incorporated by integrating over all possible phylogenetic trees and other parameters jointly. From these sampled character histories the maximum a posteriori character history can be summarized in a number of ways. The approach presented here is to calculate the marginal probabilities of character states for every small time interval along each branch, however one could also calculate the joint posterior probability of an entire character history.

During Algorithm (1) the rate-limiting step is writing conditional likelihood vectors for every small time interval along every branch on the tree, particularly when the state space of the model is large. The time required is of order *O*(*n* × *m* × *r*), where *n* is the number of taxa in the tree, *m* is the number of character states, and *r* is the number of time intervals. This is reduced by only storing conditional likelihood vectors for all time intervals during the MCMC iterations that are sampled. During unsampled (*i.e*., thinned) MCMC iterations the likelihood is calculated in the standard way storing conditional likelihood vectors only at the nodes, thus the use of the stochastic mapping algorithm has little impact on the overall computation time.

### 2.5 Onagraceae Phylogenetic Analyses

DNA sequences for Onagraceae and Lythraceae were mined from GenBank using `SUMAC` (Freyman 2015). Lythraceae was selected as an outgroup since previous molecular phylogenetic analyses place it sister to Onagraceae (Sytsma et al. 2004). In total 8 gene regions were used (7 chloroplast loci plus the nuclear ribosomal internal transcribed spacer region) representing a total of 340 taxa (292 Onagraceae taxa and 48 Lythraceae taxa). Information about the alignments and GenBank accessions used can be found in the Supporting Information Section S1.1. Phylogeny and divergence times were inferred using `RevBayes` (Höhna et al. 2016). Node ages were calibrated using five fossil calibrations and one secondary calibration. Details regarding the calibrations, the models of molecular evolution, and MCMC analyses are given in the Supporting Information Section S1.1.

### 2.6 Analyses of Mating System Evolution

The mating systems of Onagraceae species were scored as either SC or SI following Wagner et al. (2007). Most of the SC/SI assignments in Wagner et al. (2007) come from detailed family-level surveys such as Raven (1979), in which the outcrossing/selfing modes of 283 Onagraceae species were examined, and Heslop-Harrison (1990), in which compatibility tests of 48 Onagraceae species were performed. Other SC/SI assignments come from the many genus- and section-level studies cited in citetwagner2007revised such as Lewis and Lewis (1955), Plitmann et al. (1973), and Seavey et al. (1977).

For the analysis of mating system evolution, all outgroup (Lythraceae) lineages were pruned off our phylogeny, leaving 292 Onagraceae species. The species sampling fraction of extant Onagraceae species was thus *ρ* = 292/650 = 0.45, which is the number of species sampled divided by the approximate total number of Onagraceae species reported in Wagner et al. (2007). We use this sampling fraction of extant Onagraceae as the uniform taxon sampling probability *ρ*, assuming that missing species are uniformly distributed over the phylogeny (Nee et al. 1994; Yang and Rannala 1997; Höhna et al. 2011; Höhna 2014). We assumed there was no state-dependent sampling bias since we lacked complete SC/SI assignments for all Onagraceae taxa that would indicate such a bias. Finally, we accounted for uncertainty in the phylogeny and divergence times by sampling 200 trees from the posterior distribution of trees.

#### 2.6.1 HiSSE Model

To test whether diversification rate heterogeneity is associated with shifts in mating system or changes in other unmeasured traits, we used a model with 4 states that describes the joint evolution of mating system as well as an unobserved character with hidden states *a* and *b* (Figure 5). For each of the 4 states we estimated speciation (λ) and extinction (*μ*) rates. For details on priors used and the MCMC analyses see the Supporting Information Section S2.1.

The system of SI found in Onagraceae is S-RNase-based gametophytic SI (Raven 1979; Franklin et al. 1995; Igić et al. 2008). This system of SI evolved once in the common ancestor of the Asteridae and Rosidae (Steinbachs and Holsinger 2002; Igić et al. 2008; Vieira et al. 2008; Niu et al. 2017; Ramanauskas and Igić 2017), the clade that contains Onagraceae. Raven (1979) writes the system of SI in Onagraceae “is gametophytic, and involves a series of S-a1leles, with inhibition of pollen-tube growth normally in the surface layers of the stigma.” Furthermore, Raven writes that SI “seems to have been characteristic of the original common ancestor of Onagraceae, judged by the occurrence of self-incompatibility in four of the seven tribes of the family. There is no evidence for the evolution of self-incompatibility within the family once it has been lost.” Following Raven (1979), we used an irreversible model that only allowed transitions from SI to SC. However, to test the assumption of irreversibility on our results, we additionally used a model that allowed for the possibility of secondary gains of SI by permitting both transitions from SI to SC and transitions from SC to SI (see Supporting Information Section S2.3).

**Figure 5:**
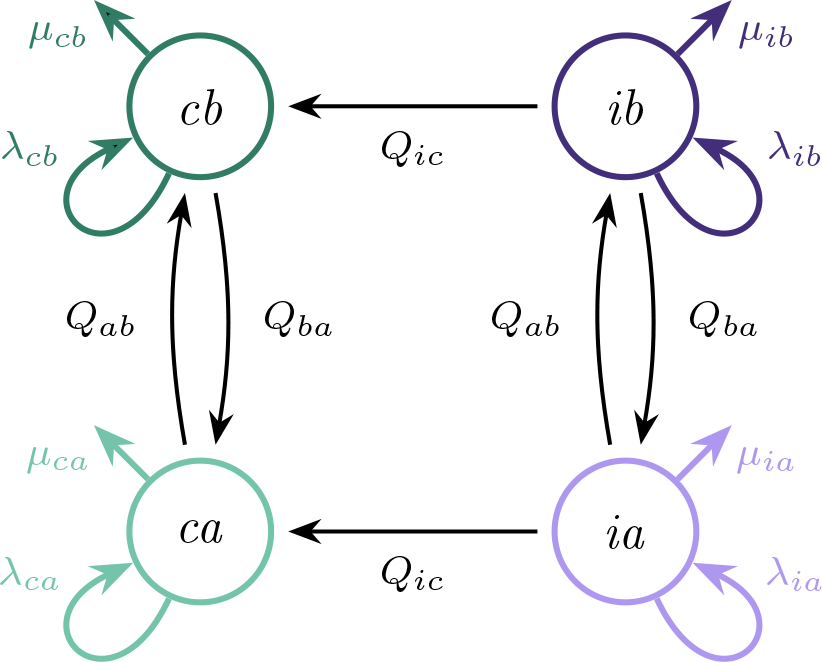
SSE model depicting states and rate parameters used to infer mating system evolution. The states are labeled *ca*, *cb*, *ia*, and *ib*, representing self-compatible hidden state *a*, self-compatible hidden state *b*, self-incompatible hidden state *a*, and self-incompatible hidden state *b*, respectively. Independent extinction *μ* and speciation λ rates were estimated for each of the 4 states, as well as the rate of transitioning from self-incompatible to self-compatible *Q_ic_* and the rates of transitioning between the hidden states *Q_ab_* and *Q_ba_*.

#### 2.6.2 Model Comparisons, Incomplete Sampling, and Error Rates

To test whether diversification-rate heterogeneity was *not* associated with shifts in mating system, we calculated a Bayes factor (Kass and Raftery 1995) to compare the mating system-dependent diversification model described above with a mating system-independent diversification model. The independent model had 4 states and the same parameters as the dependent model, except that the speciation and extinction rates were fixed so they only varied between the hidden states *a* and *b*. Hence, λ_*ca*_ was fixed to equal λ_*ia*_, λ_*cb*_ was fixed to λ_*ib*_, *μ_ca_* was fixed to *μ_ia_* and *μ_cb_* was fixed to *μ*_ib_.

To evaluate the false positive error rate and the effect of incomplete taxon sampling, we performed a series of simulations that tested the power of our models to reject false associations between shifts in mating system and diversification rate shifts. Trees were simulated under a BiSSE model, and then diversification *independent* binary characters representing mating system were simulated over the trees. To test the effect of missing data on our power to detect state-dependent diversification, the simulated datasets were pruned to have the same proportion of taxon sampling as our empirical Onagraceae dataset (45%; see Supporting Information). For each simulation replicate, Bayes factors were calculated to compare the fit of the mating system-dependent diversification model and mating system-independent diversification model. Details on the simulations are provided in the Supporting Information Section S3.1.

All Bayes factors were calculated using the stepping stone method (Xie et al. 2010; Höhna et al. 2017), as implemented in `RevBayes`. Marginal likelihood estimates were run for 50 path steps and 19000 generations within each step. The Bayes factor was then calculated as twice the difference in the natural log marginal likelihoods (Kass and Raftery 1995).

## 3 Results

### 3.1 Onagraceae Phylogeny

In our estimated phylogeny, all currently recognized Onagraceae genera (Wagner et al. 2007) were strongly supported to be monophyletic with posterior probabilities > 0.98. The crown age of Onagraceae was estimated to be 98.8 Ma (94.0 Ma - 107.3 Ma 95% HPD; Figure 6), and a summary of the divergence times of major clades within Onagraceae can be found in Supporting Information Table 3.

### 3.2 Stochastic Character Maps

Since the results from the analysis allowing for secondary gains of SI were essentially identical to the results from the analysis that assumed irreversibility and disallowed secondary gains of SI, we report here only the results from the irreversible analysis. See Supporting Information Section S2.3 for results of the analysis allowing for secondary gains.

Under the state-dependent diversification model, repeated independent losses of SI across the Onagraceae phylogeny were found to be associated with shifts in diversification rates (Figure 6). Additionally, transitions between the unobserved character states *a* and *b* were also associated with diversification rate heterogeneity. Uncertainty in the timing of diversification-rate shifts and character state transitions was generally low, but increased along long branches where there was relatively little information regarding the exact timing of transitions (Figure 7). Following the loss of SI, there was an evolutionary time lag (mean 1.97 My) until net-diversification (speciation minus extinction) turned negative (Figure 8). Since SC hidden state *b* was estimated to have positive net-diversification and SC hidden state *a* was estimated to have negative net-diversification, we calculated the time lag from the loss of SI until an evolutionary decline as the time spent following the loss of SI in hidden state *b* until transitioning to hidden state *a*. In many cases the loss of SI occurred in an ancestral lineage with positive net-diversification (hidden state *b*) followed by multiple shifts to negative net-diversification (hidden state *a*) in descendant lineages. To account for these non-independent time lags and avoid double counting the time the ancestral lineage spent in SC state *b*, we average over all (partially) dependent events. For example, the ancestral lineage spent time *t_a_* in state *b* and the left and right descendant lineages spent time *t_l_* and *t_r_* in state *b* before switching to state *a* respectively. Then, we counted the time as 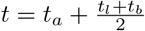.

### 3.3 Diversification Rate Estimates

Within either hidden state (*a* or *b*) SC lineages had generally higher speciation and extinction rates compared to SI lineages (Table 1 and Figure 6). Despite higher speciation and extinction rates, SC lineages had lower net-diversification compared to SI lineages. Net-diversification was found to be negative for most but not all extant SC lineages.

**Table 1:** Posterior parameter estimates of the HiSSE mating system evolution model depicted in Figure 5.

| Parameter | $X$ | Mating System | Hidden State | Mean | 95% HPD Interval |
| --- | --- | --- | --- | --- | --- |
| Speciation | $\lambda_{ca}$ | SC | $a$ | 0.12 | 0.02 – 0.23 |
| | $\lambda_{ia}$ | SI | $a$ | 0.16 | 0.09 – 0.24 |
| | $\lambda_{cb}$ | SC | $b$ | 1.66 | 0.98 – 2.41 |
| | $\lambda_{ib}$ | SI | $b$ | 0.65 | 0.45 – 0.85 |
| Extinction | $\mu_{ca}$ | SC | $a$ | 0.35 | 0.25 – 0.48 |
| | $\mu_{ia}$ | SI | $a$ | 0.04 | 0.00 – 0.09 |
| | $\mu_{cb}$ | SC | $b$ | 1.36 | 0.65 – 2.19 |
| | $\mu_{ib}$ | SI | $b$ | 0.10 | 0.00 – 0.29 |
| Net-diversification | $r_{ca}$ | SC | $a$ | -0.23 | -0.32 – -0.14 |
| | $r_{ia}$ | SI | $a$ | 0.13 | 0.05 – 0.19 |
| | $r_{cb}$ | SC | $b$ | 0.30 | 0.15 – 0.46 |
| | $r_{ib}$ | SI | $b$ | 0.55 | 0.39 – 0.71 |
| Transition | $Q_{ic}$ | SI → SC | $a/b$ | 0.22 | 0.16 – 0.28 |
| | $Q_{ab}$ | SI/SC | $a \rightarrow b$ | 0.01 | 0.00 – 0.02 |
| | $Q_{ba}$ | SI/SC | $b \rightarrow a$ | 0.33 | 0.16 – 0.52 |

#### 3.3.1 Model Comparisons, Incomplete Sampling, and Error Rates

For the Onagraceae dataset, the state-dependent diversification model of mating system evolution (Figure 5) was “decisively” supported over the state-independent diversification model with a Bayes factor (2*ln*BF) of 19.9 (Jeffreys 1961). Bayes factors calculated using simulated datasets showed that the false positive error rate was low even despite the poor taxon sampling present in our empirical dataset (Figure 9). The false positive rate for “strong” support (2*ln*BF > 6; Kass and Raftery 1995) was 0.05, and the false positive rate for “very strong” support (2*ln*BF > 10; Kass and Raftery 1995) was 0.0.

**Figure 6:**
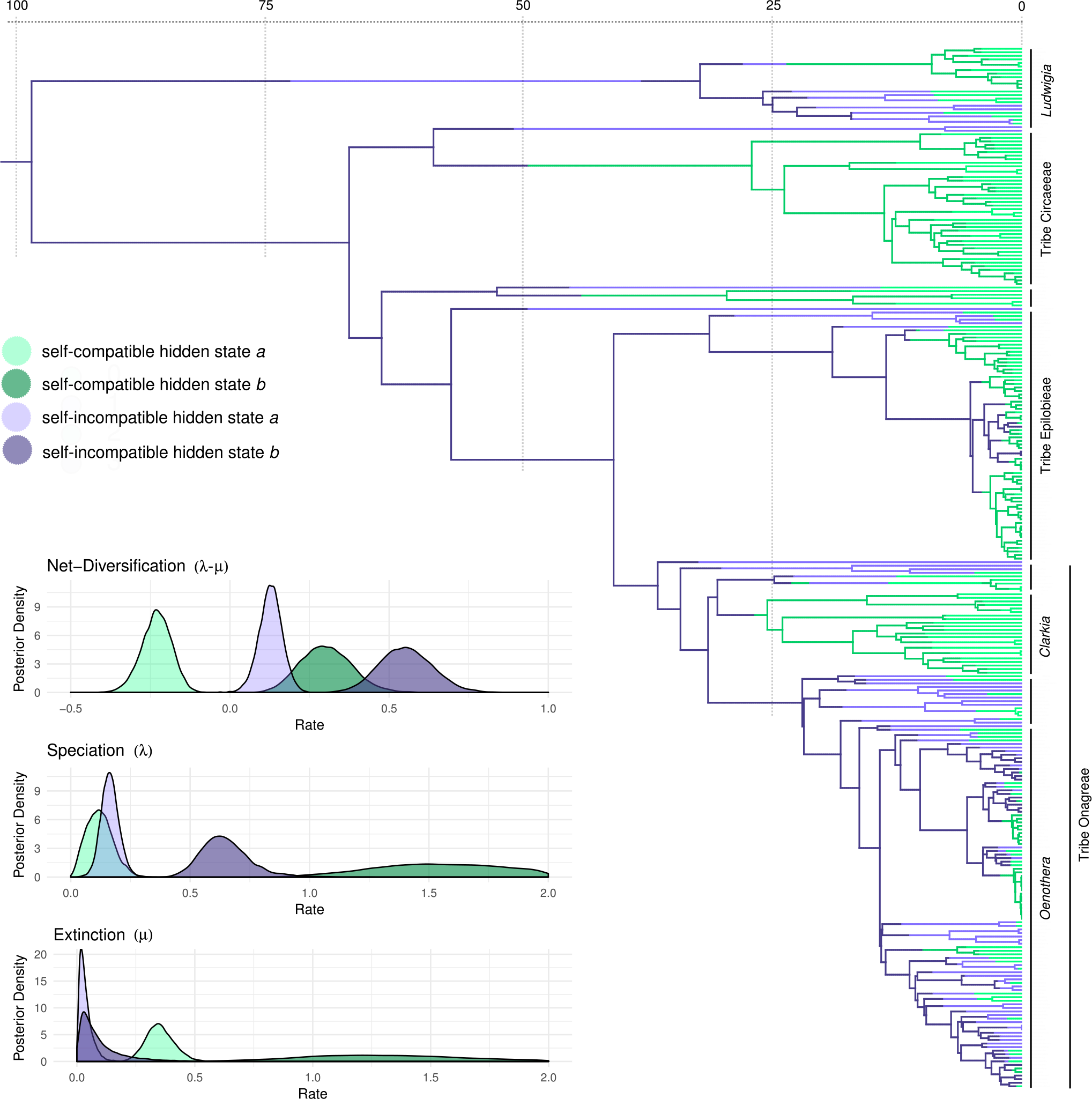
Maximum a posteriori reconstruction of mating system evolution and shifts in diversification rates in Onagraceae. Divergence times in millions of years are indicated by the axis at the top. Note that in this marginal summary reconstruction some transitions are displayed such as the loss and regain of self-incompatibility that were impossible in any single sampled character history. This indicates high uncertainty in the exact timing of transitions (see Figure 7). The inset panels show posterior densities of net-diversification (λ − *μ*), speciation (λ), and extinction (*μ*) rates in millions of years. Changes in mating system and an unobserved character (hidden states *a* and *b*) are both associated with diversification rate heterogeneity. Within either hidden state (*a* or *b*) self-compatible lineages have higher extinction and speciation rates yet lower net-diversification rates compared to self-incompatible lineages.

**Figure 7:**
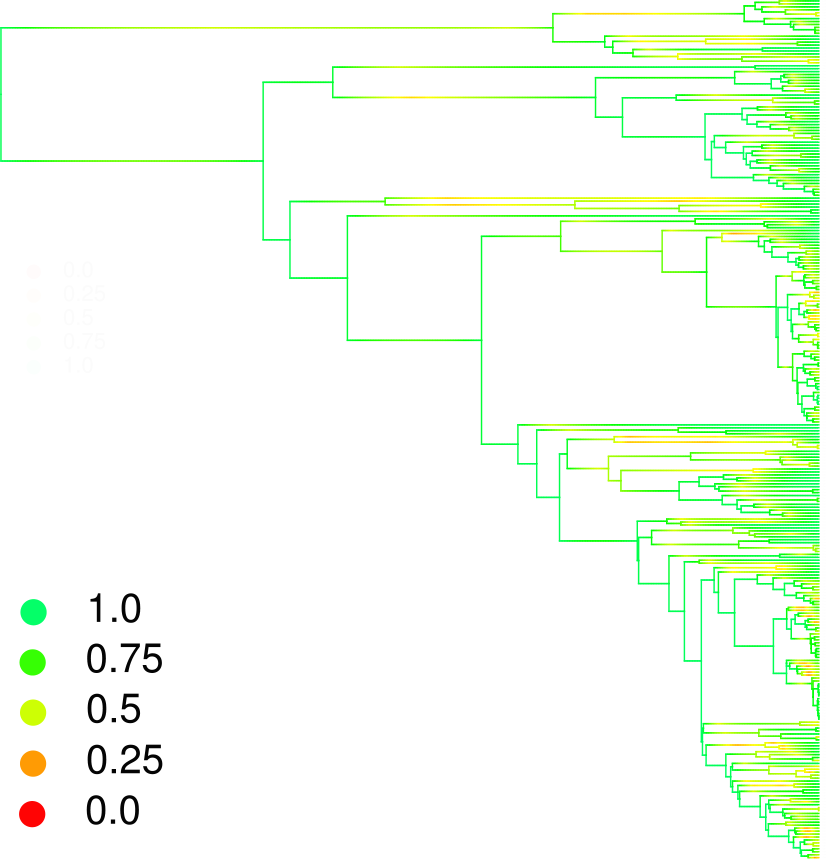
Posterior probabilities of the maximum a posteriori reconstruction of mating system evolution and shifts in diversification rates in Onagraceae. Marginal posterior probabilities of the character states shown in Figure 6. Uncertainty was highest along long branches where there was relatively little information regarding the timing of transitions.

**Figure 8:**
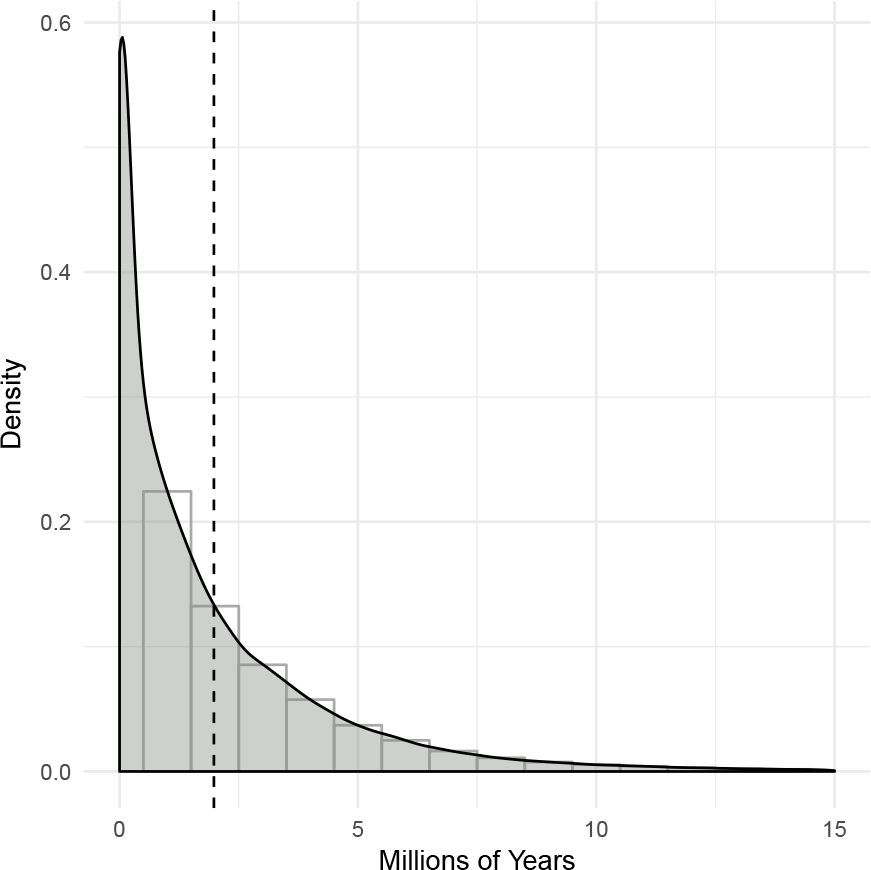
The time lag from the loss of self incompatibility until the onset of evolutionary decline. The time in millions of years after the loss of self-incompatibility until the net-diversification rate became negative measured over 10000 stochastic character map samples. The mean time lag until evolutionary decline was 1.97 million years (indicated by a dashed line).

**Figure 9:**
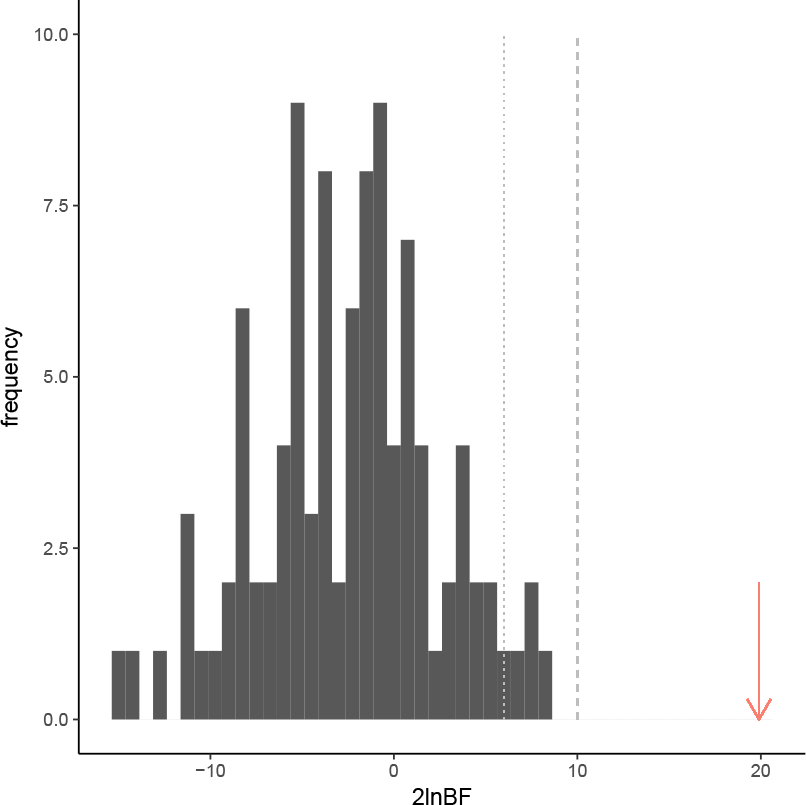
Bayes factors (2*ln*BF) comparing the fit of the state-dependent diversification model of mating system evolution with the state-independent diversification model. The red arrow indicates the “decisive” support found for the empirical Onagraceae data (2*ln*BF = 19.9; Jeffreys 1961). The dark grey bars represent Bayes factors calculated for 100 datasets simulated under a state-independent diversification model and pruned to have the same proportion of missing species as the empirical Onagraceae dataset. The dotted light grey line indicates “strong” support (2*ln*BF > 6; Kass and Raftery 1995), and the dashed light grey line indicates “very strong” support (2*ln*BF > 10; Kass and Raftery 1995). Even with the poor taxon sampling present in the Onagraceae dataset, our power to reject false positives was high.

## 4 Discussion and Conclusion

The stochastic character map results reveal that the loss of SI has different short term and long term macroevolutionary consequences. Lineages with relatively recent losses of SI like *Epilobium* are undergoing a burst in both speciation and extinction rates with a positive net-diversification rate. However, lineages that have long been SC such as *Fuchsia* (Tribe Circaeeae) and *Clarkia* are in a previously unrecognized evolutionary decline. These lineages went through an increase in both speciation and extinction rates a long time ago—after the loss of SI— but now only the extinction rates remain elevated and the speciation rates have declined, resulting in a negative net-diversification rate. The time lag until this evolutionary decline was measured as the time spent following the loss of SI in hidden state *b* (positive net-diversification) until transitioning to hidden state *a* (negative net-diversification). By mapping the time spent in each hidden state, the stochastic character maps quantified the speed of the evolutionary decline in SC lineages. These results are robust to phylogenetic uncertainty (by averaging over a posterior distribution of trees), to assumptions of mating system irreversiblity (Supporting Information for results allowing for secondary gains of SI), and to the effect of missing species sampling (false positive error rate calculated using simulations).

While the mean time from the loss of SI until evolutionary decline was 1.97 My, there was a large amount of variation in time estimates (Figure 8). This variation could be due to differences in the realized selfing/outcrossing rates of different SC lineages. Lineages with higher selfing rates likely build up load due to weakly deleterious mutations more quickly, leading to a more rapid mutational meltdown and eventual evolutionary decline (Lynch et al. 1995a,b; Wright et al. 2008). Furthermore, even if mutational load is low, the loss of genetic variation in highly selfing lineages will reduce the probability that such lineages can respond adequately to natural selection, such as imposed by a changing or new environment, thus increasing the potential for extinction. SC lineages with high outcrossing rates and less inbreeding, on the other hand, likely have larger effective population sizes and lower genetic load (Wright et al. 2008), thus delaying the onset of higher extinction rates. A limitation of our analysis is that our species data was coded simply SI or SC, a more nuanced exploration of the macroevolutionary impact of selfing would require hard to measure selfing/outcrossing rates from a large number of species across the phylogeny.

These results confirm theory about the macroevolutionary consequences of selfing (Stebbins 1957; Grant 1981). These consequences include the increased probability of going extinct due to the accumulation of harmful mutations (Lynch et al. 1995a,b; Wright et al. 2008) and an increased rate of speciation which may be driven by higher among-population differentiation and reproductive assurance that facilitates colonization of new habitats (Baker 1955; Hartfield 2016). The advantages of reproductive assurance may explain why transitions to SC occur repeatedly (Igić et al. 2008; Lande and Schemske 1985). However, our results reveal that this advantage in Onagraceae is short-lived; the burst of increased speciation following the loss of SI eventually declines, possibly due to failing to adapt to changing conditions and the accumulation of deleterious mutations. The overall macroevolutionary pattern is one in which SC Onagraceae lineages undergo rapid bursts of increased speciation that eventually decline, doomed by intensified extinction and thus supporting Stebbins’ hypothesis of selfing as an evolutionary dead-end (Stebbins 1957). These results provide empirical evidence for the “senescing” diversification rates predicted in highly selfing lineages by Ho and Agrawal (2017), who proposed that primarily selfing lineages may at first diversify at higher rates than outcrossing lineages but over time slow down due to elevated extinction rates. Similar results were previously found in Primulaceae by de Vos et al. (2014), where SC non-heterostylous lineages were found to “live fast and die young” compared to SI heterostylous lineages.

Our findings corroborate previous analyses performed in the plant families Solanceae (Goldberg et al. 2010), Primulaceae (de Vos et al. 2014), and Orchidaceae (Gamisch et al. 2015) where SC lineages were also found to have lower net-diversification rates than SI lineages. Our results, however, are the first to use a HiSSE model to show that this pattern is supported even when other unmeasured factors affect diversification rate heterogeneity. Intuitively, it is clear that no single factor drives all diversfication rate heterogeneity in diverse and complex clades such as Onagraceae. Indeed, in some lineages of *Oenothera* the loss of sexual recombination and segregation due to extensive chromosome translocations (a condition called Permanent Translocation Heterozygosity) is associated with increased diversification rates (Johnson et al. 2011). Furthermore, other factors such as polyploidy and shifts in habitat, growth form, or life cycle may impact diversification rates (Mayrose et al. 2011; Donoghue 2005; Eriksson and Bremer 1992). Interpretating the hidden states of an SSE model can be challenging (Caetano et al. 2018). Depending on the diversification rates estimated there were different but equally valid ways to make sense of the hidden states in our analysis: (1) if the diversification rates varied between SC and SI, but not between hidden states *a* and *b*, we could conclude that shifts in mating system explained all diversification rate heterogeneity; (2) if the diversification rates did not vary between SC and SI, but did vary between hidden states *a* and *b* we could conclude that there were background rate changes unassociated with mating system and that mating system evolution was not associated with rate shifts; or (3) if the diversification rates varied both between SC/SI and between hidden states *a/b*, then depending on the phylogenetic pattern of the hidden states they could represent the different long and short term consequences of the loss of SI. Our results are congruent with this last interpretation, and we interpret the phylogenetic pattern of the hidden states to represent the temporal decay of diversification rates in SC lineages. It is important to note, however, that our HiSSE-based analysis allowed for any of those three outcomes unlike BiSSE-based analyses.

Stochastic character mapping of state-dependent diversification, can be a powerful tool for examining the timing and nature of both shifts in diversification rates and character state transitions on a phylogeny. Character mapping reveals which stages of the unobserved character a lineage goes through; e.g. after the loss of self-incompatibility transitions are predominantly from hidden state *b* to *a*, representing shifts from positive net-diversification to negative net-diversification. Furthermore, character mapping infers the state of the lineages in the present and so reveals which tips of the phylogeny are currently undergoing positive or negative net-diversification. If stochastic character mapping is used with an SSE model in which some or even all states are hidden (no observed states), then our method will “paint” the location of shifts in diversification rate regimes over the tree. Distributions of character map samples could be used for posterior predictive assessments of model fit (Nielsen 2002; Bollback 2006; Höhna et al. 2018) and for testing whether multiple characters coevolve (Huelsenbeck et al. 2003; Bollback 2006). Our hope is that these approaches enable researchers to examine the macroevolutionary impacts of the diverse processes shaping the tree of life with increasing quantitative rigor.

## 5 Acknowledgements

Thank you to Bruce Baldwin, John Huelsenbeck, Emma Goldberg, Michael Landis, Seema Sheth, and Carl Rothfels and his lab group for discussions that have improved our work. W.A.F. was supported by grants from the National Science Foundation (GRFP DGE 1106400, DDIG DEB 1601402, and DEB 1655478). Computations were performed on the Savio computational cluster provided by the Berkeley Research Computing program at the University of California, Berkeley.

## S1 Onagraceae Phylogenetic Analyses

### S1.1 Methods

#### S1.1.1 Supermatrix Assembly

DNA sequences for Onagraceae and Lythraceae were mined from GenBank using `SUMAC` (Freyman 2015). Lythraceae was selected as an outgroup since previous molecular phylogenetic analyses place it sister to Onagraceae (Sytsma et al. 2004). `SUMAC` assembled an 8 gene supermatrix (7 chloroplast loci plus the nuclear ribosomal internal transcribed spacer region) representing a total of 340 taxa. Table S1 summarizes the genes used, their length, and the percent of missing data. We counted N’s in the downloaded sequences as missing data, but we did not count gaps introduced by the alignment algorithm as missing data. Sequences were aligned using `MAFFT v7.123b` (Katoh and Standley 2013). The default settings in `MAFFT` were used except that proper sequence polarity was ensured by using the direction adjustment option. Alignments were then concatenated resulting in chimeric operational taxonomic units (OTUs) that do not necessarily represent a single individual.

**Table S1.** DNA regions mined from GenBank. A total of 340 taxa were included.

| DNA Region | # Taxa | Aligned Length | # Variable Sites | Missing data (%) | Taxon Coverage Density |
| --- | --- | --- | --- | --- | --- |
| ITS | 250 | 904 | 481 | 26.5 | 0.735 |
| matK | 42 | 895 | 276 | 87.6 | 0.124 |
| ndhF | 39 | 1085 | 429 | 88.5 | 0.115 |
| pgiC | 66 | 7664 | 3828 | 80.6 | 0.194 |
| rbcL | 108 | 1427 | 388 | 68.2 | 0.318 |
| rpl16 | 54 | 1139 | 343 | 84.1 | 0.159 |
| rps16 | 78 | 1056 | 335 | 77.1 | 0.229 |
| trnL-trnF | 261 | 1434 | 644 | 23.2 | 0.768 |

**Table S2.** Fossil and secondary calibrations used as priors in the Bayesian divergence time analysis. Units are in millions of years.

| Group | Calibration Type | Placement | Prior Distribution | Mean | SD | Offset | Reference |
| --- | --- | --- | --- | --- | --- | --- | --- |
| *Circaea* | fossil | stem | lognormal | 10 | 2 | 12 | (Grímsson et al. 2012) |
| Tribe Epilobieae | fossil | stem | lognormal | 10 | 2 | 12 | (Grímsson et al. 2012) |
| *Fuchsia* section *Skinnera* | fossil | stem | lognormal | 10 | 2 | 23 | (Lee et al. 2013) |
| Lythraceae | fossil | crown | lognormal | 20 | 2 | 81.5 | (Graham 2013) |
| *Ludwigia* | fossil | stem | lognormal | 10 | 2 | 57.6 | (Zhi-Chen et al. 2004) |
| Onagraceae + Lythraceae | secondary | crown | normal | 93 | 5 | 0 | (Sytsma et al. 2004) |

#### S1.1.2 Phylogenetic Analyses

Divergence times and phylogeny were jointly estimated using `RevBayes` (Höhna et al. 2014, 2016). Estimates were time calibrated using six node calibrations: four stem fossil calibrations, one crown fossil calibration, and a secondary calibration for the root split between Onagraceae and Lythraceae (Table S2). An uncorrelated lognormal relaxed clock model was used, and each of the eight gene partitions were assigned independent GTR substitution models (Tavaré 1986; Rodriguez et al. 1990). Rate variation across sites was modeled under a gamma distribution approximated by four discrete rate categories (Yang 1994). The constant rate birth-death-sampling tree prior (Nee et al. 1994; Yang and Rannala 1997) was used with the probability of sampling species at the present (ρ) set to 0.27. *ρ* was calculated by dividing the number of extant species sampled in the supermatrix (340) by the sum of the number of species recognized in Onagraceae (~650) and in Lythraceae (~620).

Four independent MCMC analyses were performed. Each MCMC ran for 15000 generations, where each generation consisted of 837 randomly scheduled Metropolis-Hastings moves. This resulted in four chains that each performed a total of 12,555,000 MCMC steps. Samples of the posterior distribution were drawn every 10 generations, and the first 50% of samples from each chain were discarded as burnin resulting in 750 trees sampled from each of the 4 independent chains. Convergence was assessed by ensuring the effective sample size of each parameter was over 200 for each independent chain. The maximum a posteriori (MAP) tree was then calculated from the combined 3000 tree samples of all 4 chains.

### S1.2 Results

All Onagraceae genera described in Wagner et al. (2007) were recovered as monophyletic in the MAP summary tree with posterior probabilities > 0.95 (Figure S4). Onagraceae was found to diverge from Lythraceae at 111.3 My (95% HPD interval 106.0 - 116.6 My). Divergence time estimates of other major clades and 95% HPD intervals can be seen in Table S3.

**Table S3.** Divergence time estimates of major clades.

| Clade | Age Type | Mean Age (Ma) | 95% HPD Min | 95% HPD Max |
| --- | --- | --- | --- | --- |
| Onagraceae + Lythraceae | crown | 111.3 | 106.0 | 116.6 |
| Onagraceae | crown | 98.8 | 94.0 | 107.3 |
| *Ludwigia* | crown | 32.1 | 31.3 | 50.6 |
| Tribe Circaeae | stem | 58.7 | 42.3 | 65.0 |
| Tribe Circaeae | crown | 27.0 | 24.6 | 29.8 |
| *Fuchshia* | crown | 23.7 | 23.0 | 24.1 |
| *Lopezia* | crown | 29.5 | 25.3 | 36.7 |
| Tribe Epilobieae | stem | 40.8 | 39.3 | 47.0 |
| Tribe Epilobieae | crown | 31.2 | 31.2 | 40.6 |
| *Chamerion* | crown | 14.9 | 12.6 | 25.2 |
| *Epilobium* | crown | 18.9 | 15.7 | 22.9 |
| Tribe Onagreae | stem | 40.8 | 39.3 | 47.0 |
| Tribe Onagreae | crown | 36.4 | 31.6 | 40.5 |
| *Taraxia* | crown | 17.0 | 9.9 | 19.5 |
| *Gayophytum* | crown | 3.1 | 1.8 | 6.1 |
| *Clarkia* | crown | 25.4 | 24.6 | 31.2 |
| *Eremothera* | crown | 10.4 | 8.1 | 15.1 |
| *Camissonia* | crown | 9.7 | 6.4 | 14.6 |
| *Eulobus* | crown | 4.7 | 2.6 | 8.1 |
| *Chylismia* | crown | 14.4 | 10.8 | 19.0 |
| *Oenothera* | crown | 14.2 | 13.0 | 17.6 |

**Figure S1:**
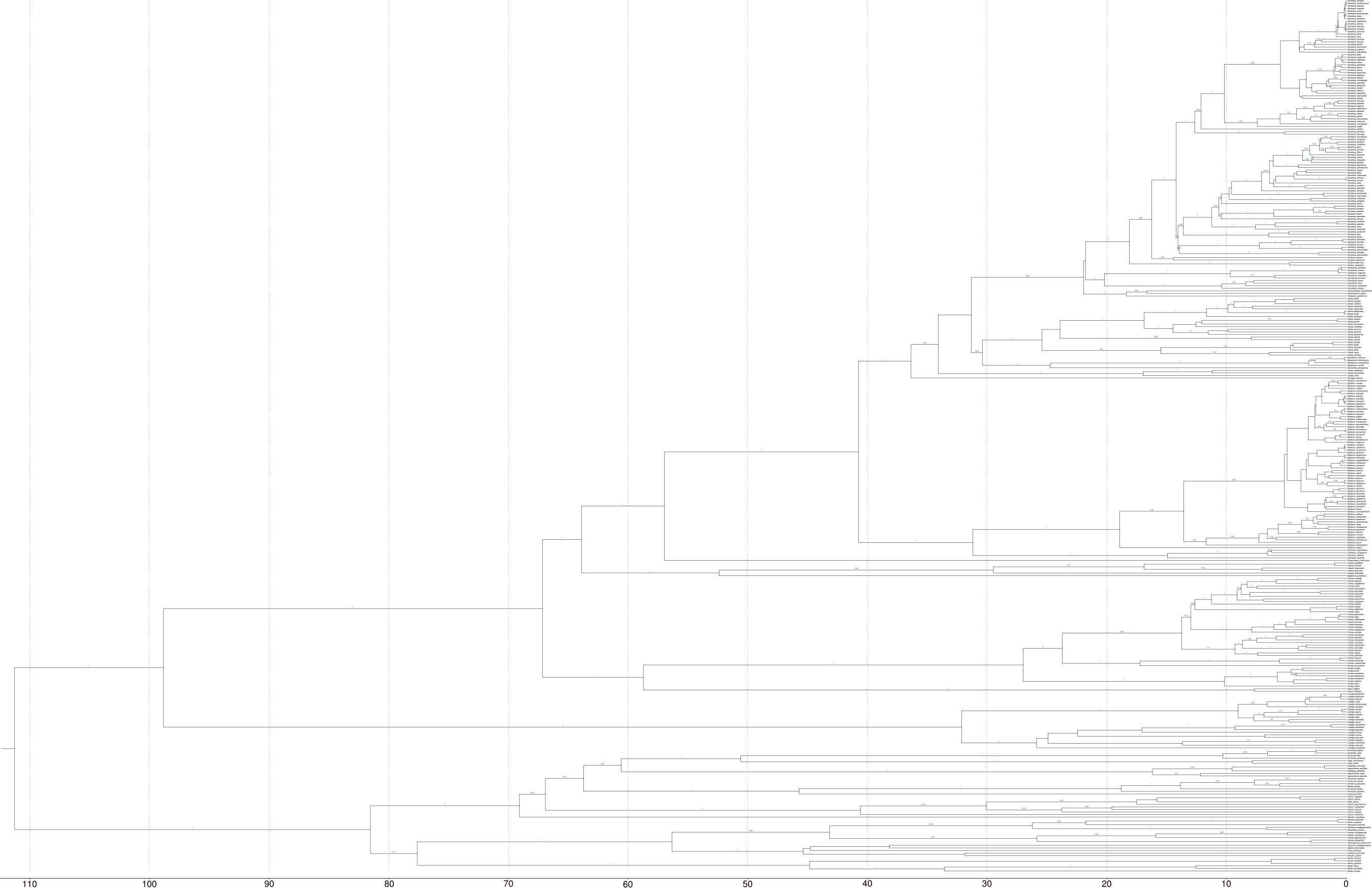
Maximum a posteriori estimate of phylogeny and divergence times of Onagraceae. Divergence times in millions of years are indicated by the axis at the bottom. Posterior probabilities > 0.50 are labeled on the branches.

## S2 Mating System Evolution Analyses

### S2.1 Model Priors

Model parameter priors are listed in Table S4. The rate of loss of self-incompatibility (*q_ic_*), and the rates of switching between hidden states *a* and *b* (*q_ab_* and *q_ba_*) were each given an exponential distribution with a mean of *n*/Ψ_*l*_, where Ψ_*l*_ is the length of the tree Ψ and *n* is the expected number of transitions. *n* was given an exponential hyperprior with a mean of 20.

The speciation and extinction rates were drawn from exponential priors with a mean equal to an estimate of the net diversification rate *d̂*. Under a constant rate birth-death process not conditioning on survival of the process, the expected number of lineages at time *t* is given by:

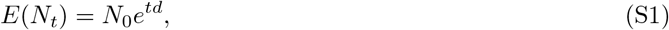

where *N*_0_ is the number of lineages at time 0 and *d* is the net diversification rate λ − *μ* (Nee et al. 1994; Höhna 2015). Therefore, we estimate *d̂* as:

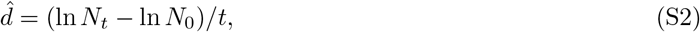

where *N_t_* is the number of lineages in the clade that survived to the present, *t* is the age of the root, and *N*_0_ = 2. The root state probabilities π were set to start the process equally in either self-incompatible hidden state *a* or self-incompatible hidden state *b*.

**Table S4:** Model parameter names and prior distributions. See the main text for complete description of model parameters and prior distributions. Ψ_*l*_ represents the length of tree Ψ and *d̂* is the expected diversification rate under a constant rate birth-death process.

| Parameter | $X$ | $f(X)$ |
| --- | --- | --- |
| Speciation self-incompatible $a$ | $\lambda_{ia}$ | Exponential($\lambda = 1/\hat{d}$) |
| Speciation self-incompatible $b$ | $\lambda_{ib}$ | Exponential($\lambda = 1/\hat{d}$) |
| Speciation self-compatible $a$ | $\lambda_{ca}$ | Exponential($\lambda = 1/\hat{d}$) |
| Speciation self-compatible $b$ | $\lambda_{cb}$ | Exponential($\lambda = 1/\hat{d}$) |
| Extinction self-incompatible $a$ | $\mu_{ia}$ | Exponential($\lambda = 1/\hat{d}$) |
| Extinction self-incompatible $b$ | $\mu_{ib}$ | Exponential($\lambda = 1/\hat{d}$) |
| Extinction self-compatible $a$ | $\mu_{ca}$ | Exponential($\lambda = 1/\hat{d}$) |
| Extinction self-compatible $b$ | $\mu_{cb}$ | Exponential($\lambda = 1/\hat{d}$) |
| Rate of loss of self-incompatibility | $q_{ic}$ | Exponential($\lambda = \Psi_l/n$) |
| Rate of $a \rightarrow b$ | $q_{ab}$ | Exponential($\lambda = \Psi_l/n$) |
| Rate of $b \rightarrow a$ | $q_{ba}$ | Exponential($\lambda = \Psi_l/n$) |
| Expected number of transitions | $n$ | Exponential($\lambda = 1/20$) |

### S2.2 MCMC Analyses

To account for uncertainty in phylogeny and divergence times 200 independent MCMC analyses were performed, each sampling a tree from the posterior distribution of trees generated during the phylogenetic analyses. All outgroup (Lythraceae) lineages were pruned off, leaving 292 Onagraceae species. The probability of sampling species at the present (*ρ*) was set to 292/650 = 0.45, which is the number of Onagraceae species sampled divided by the approximate number of species in Onagraceae. Each MCMC run drew 10000 samples from the posterior distribution, with 190 randomly scheduled Metropolis-Hastings moves per sample. The first 10% of samples from each run were discarded as burnin. For each run, all parameters had effective sample sizes greater than 200, and the mean effective sample size of the posterior across all 200 tree samples was 1161.6. Estimates of the diversification rates were made by combining samples from all 200 independent runs.

### S2.3 Mating System Analysis Allowing for Reversals

To test for secondary gains of self-incompatibility, we repeated the analysis described above but instead of fixing *q_ci_* = 0 we estimated both *q_ci_* and *q_ic_*. Like *q_ic_*, we assigned *q_ci_* an exponential prior with λ = Ψ_*l*_/*r*, where the expected number of reversals *r* was exponentially distributed with parameter λ = 1/20.

The results from the analysis allowing for secondary reversals were nearly identical to the results from the analysis disallowing secondary reversals (Figure S2 and S3). The transition rate from SI to SC was 0.19 (0.10 - 0.26 95% HPD), slightly lower than when reversals were disallowed. The rate of secondary reversals SC to SI was 2.52 × 10^−3^ (1.94 × 10^−5^ − 5.88 × 10^−3^ 95% HPD), essentially zero. This resulted in sampled character histories nearly identical to those shown in the main text estimated under the irreversible model.

The diversification rates estimated when allowing for reversals were very similar to the diversification rates estimated when not allowing for reversals. Within either hidden state (*a* or *b*) SC lineages had generally higher speciation and extinction rates compared to SI lineages (Figure S3). SC lineages in state *a* had a speciation rate of 0.14 (0.03 - 0.25 95% HPD) compared to 0.15 (0.08 − 0.22 95% HPD) in SI lineages in state *a*. For SC lineages in state *b* the speciation rate was 1.60 (1.00 - 2.29 95% HPD) compared to 0.61 (0.44 − 0.82 95% HPD) in SI lineages in state *b*. Similarly, SC lineages in state *a* had an extinction rate of 0.36 (0.25 - 0.48 95% HPD) compared to 0.04 (0.00 - 0.09 95% HPD) in SI lineages in state *a*. For SC lineages in state *b* the extinction rate was 1.27 (0.61 - 2.02 95% HPD) compared to 0.08 (0.00 - 0.25 95% HPD) in SI lineages in state *b*.

Despite higher speciation and extinction rates, SC lineages had lower net diversification compared to SI lineages. Net diversification was found to be negative for most but not all extant SC lineages. The net diversification rate for SC lineages in state *a* was −0.23 (−0.31 - −0.14 95% HPD), compared to 0.11 (0.04 - 0.18 95% HPD) in SI lineages in state *a*. For SC lineages in state *b* the net diversification rate was 0.33 (0.17 - 0.50 95% HPD), compared to 0.53 (0.36 - 0.70 95% HPD) in SI lineages in state *b*.

**Figure S2:**
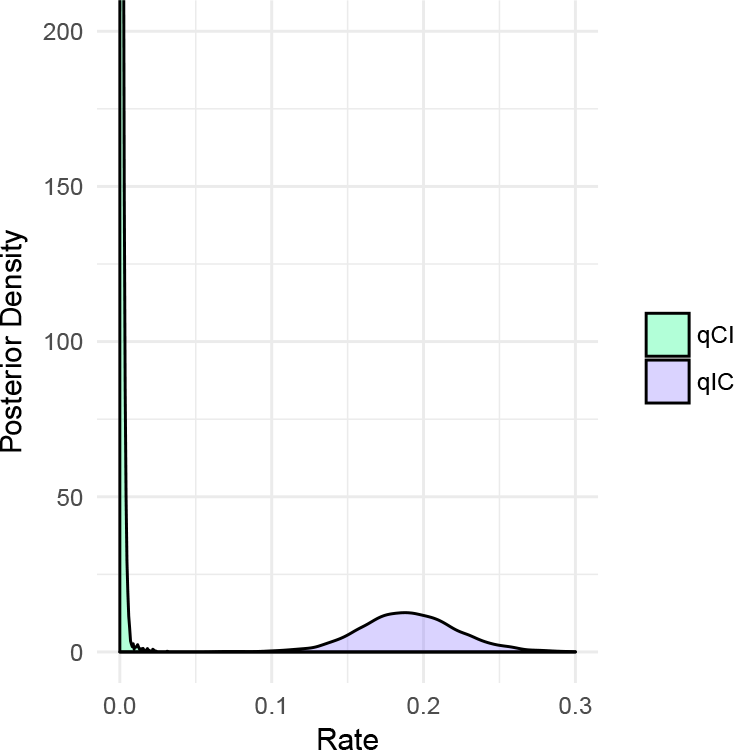
Posterior density of transition rate estimates when allowing for reversals. The estimated rate of secondary gains of SI (*Q_ci_*; green) was 2.52 × 10^−3^ (1.94 × 10^−5^ - 5.88 × 10^−3^ 95% HPD). The estimated rate of the loss of SI (*Q_ic_*; purple) was 0.19 (0.10 - 0.26 95% HPD).

## S3 Simulations

### S3.1 Simulated Datasets

100 datasets were simulated under a model where the observed binary character was diversification rate independent yet an unobserved binary character drove background diversification rate heterogeneity. To test the effect of missing data on our power to detect state-dependent diversification, we simulated datasets with the same proportion of taxon sampling as our empirical Onagraceae dataset (45%; see details above for how this number was calculated). First trees were simulated under BiSSE (Maddison et al. 2007) as implemented in the R package `diversitree` (FitzJohn 2012). The binary character represented hidden states *a* and *b* with diversification rates λ_*a*_ = 1.0, λ_b_ = 2.0, *μ*_*a*_ = 0.4, and *μ*_*b*_ = 0.1. The rate of change between hidden states *a* and *b* was set to *q_ab_* = *q_ba_* = 0.1. This resulted in trees that were qualitatively similar in shape to the empirically estimated Onagraceae tree, with a mix of early diverging depauperate clades and more rapidly radiating recent clades (Figure S4). To simulate incomplete sampling, 55% of the extant tips were randomly pruned off the tree. After pruning, tree samples were discarded unless they had between 100 and 200 sampled lineages that survived to the present. This restriction ensured that the simulated datasets were not too small for reliable inference and yet not so large to be computationally infeasible. Furthermore, we discarded datasets that had fewer than 20% of the tips in either hidden state to ensure that the trees were generated under a sufficiently heterogenous process.

**Figure S3:**
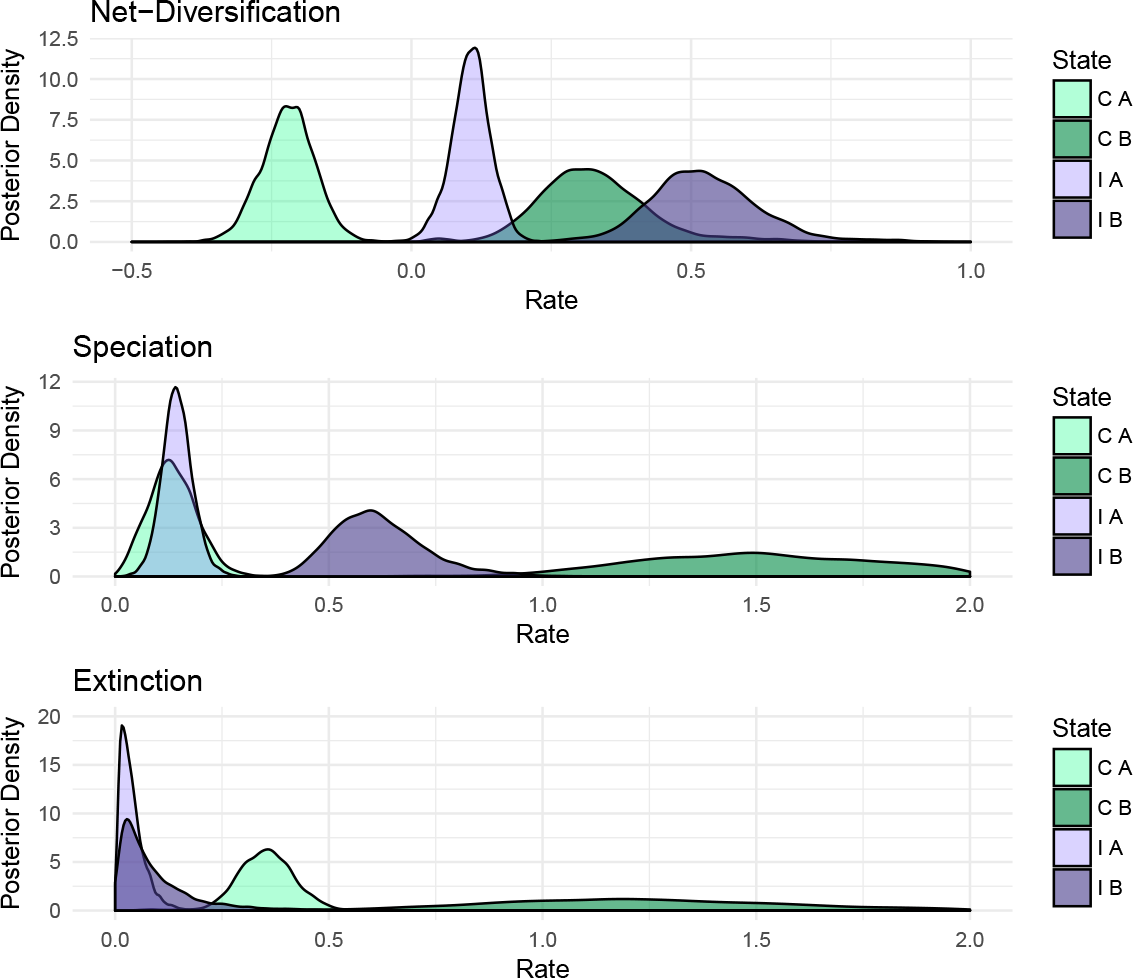
Posterior density of diversification rates when allowing for reversals. Diversification rate estimates were nearly identical to those estimated when not allowing for reversals (compare with Figure 6 in the main text).

Once the trees were simulated, diversification independent binary characters were simulated over the trees. These characters represented the observed character (mating system) and so were simulated under an irreversible model where the allowed transition occurred with the rate 10/Ψ_*s*_, where Ψ_*s*_ is the length of the simulated tree. This represents an expected 10 irreversible transitions over the length of the tree, and resulted in simulated datasets with a proportion of either state similar to the proportion of self-compatible/self-incompatible in the empirical Onagraceae dataset. These diversification independent characters were then used to calculate Bayes factors that compared the fit of the diversification dependent model to the diversification independent model of mating system. For details on how Bayes factors were calculated see the main text. The false positive error rate was calculated as the percent of simulation replicates in which the Bayes factor supported the false dependent model over the true independent model.

**Figure S4:**
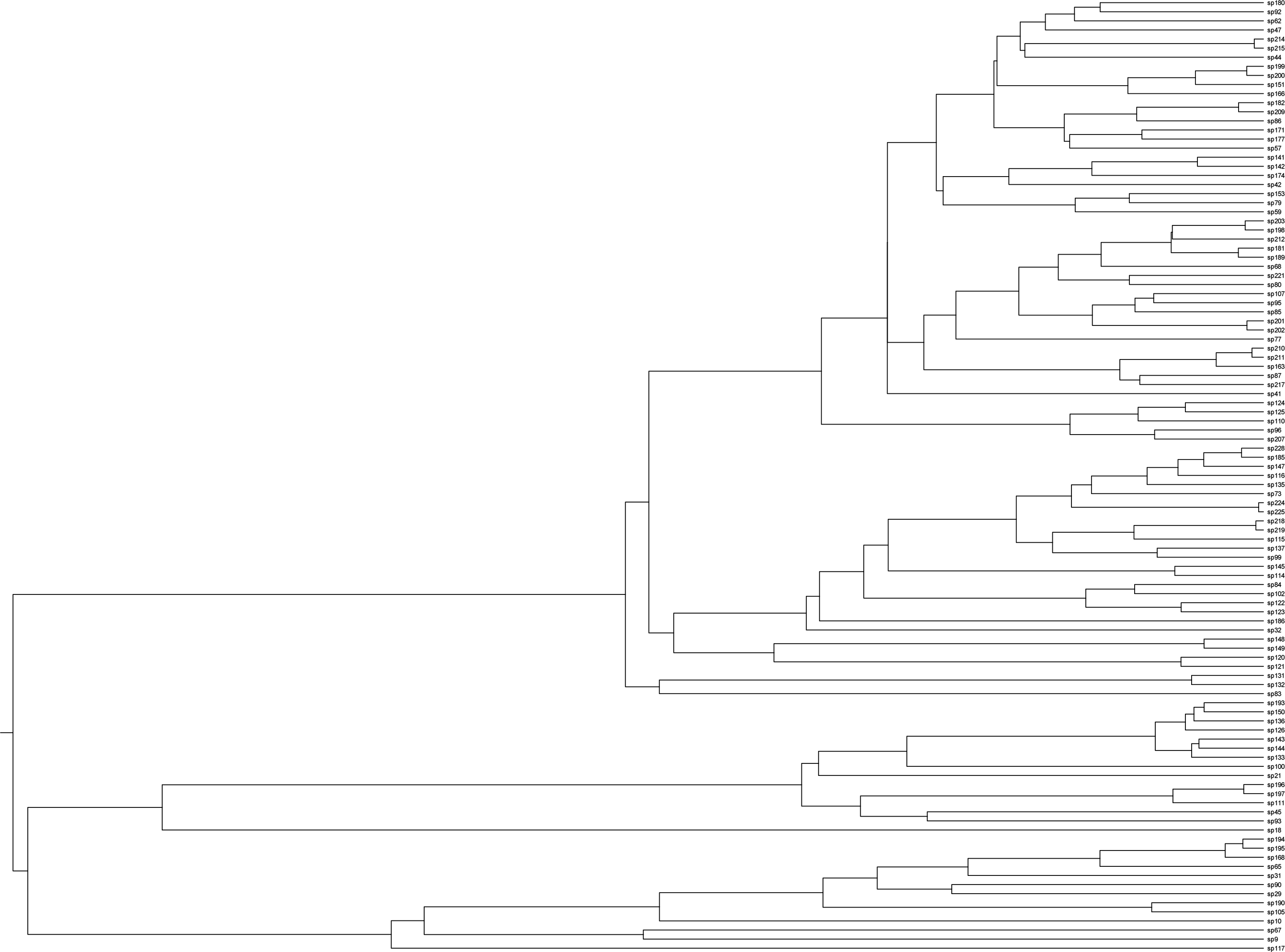
One of the trees simulated under BiSSE used to calculate the false positive error rate. Trees were simulated under a heterogenous diversification process to result in a mix of early diverging depauperate clades and more rapidly radiating recent clades. The tree is shown after 55% of the extant lineages were randomly pruned to replicate incomplete sampling in a reconstructed phylogeny.

